# Rapid growth and the evolution of complete metamorphosis in insects

**DOI:** 10.1101/2024.02.12.579885

**Authors:** Christin Manthey, C Jessica E Metcalf, Michael T Monaghan, Ulrich Karl Steiner, Jens Rolff

## Abstract

More than 60% of all animal species are insects that undergo complete metamorphosis. The key innovation of these holometabolous insects is a pupal stage between the larva and adult when most structures are completely rebuilt. Why this extreme lifestyle evolved is unclear. Here we test the hypothesis that a trade-off between growth and differentiation explains the evolution of this novelty. Using a comparative approach, we find that holometabolous insects grow much faster than hemimetabolous insects. Using a theoretical model, we then show how holometaboly evolves under a growth-differentiation trade-off and identify conditions under which such temporal decoupling of growth and differentiation is favored. Our work supports the notion that the holometabolous life history evolved to remove developmental constraints on fast growth, primarily under high mortality.

---

Metamorphosis, the life-cycle discontinuity between larval and adult phenotypes, is found in the majority of animal taxa (*1*) and is commonly explained as an adaptation that allows organisms to optimize their phenotypes to different habitats or diets (*2, 3*). More than 60% of all extant animal species are insects (*4*), 80% of which undergo an extreme form of metamorphosis: holometaboly. Holometabolous insects are monophyletic (*5*) and radiated in the Mesozoic (*6*). In holometabolous insects, the pupal stage is an evolutionarily novel phenotype interposed between the larval and the adult stages. During this phase of development, most of the body is radically reorganized, both externally and internally (*7–10*), building on embryonic tissue that has passed through the larval stages in a determined but not differentiated state (*11–13*). In the pupa, differentiation of adult organs takes place as does the replacement of almost all larval organs, leading to the emergence of a distinct adult life form (*8, 9, 14, 15*). Holometaboly has fascinated students of natural history since Aristotle (*10, 16*)’, yet the evolution of this extreme form of metamorphosis is puzzling: if decoupling different life stages is the key adaptation of metamorphosis (*1, 17*), simpler forms of metamorphosis such as hemimetaboly should suffice. What is then the extra driver for the evolution of complete metamorphosis?

One intriguing hypothesis to explain the evolution of complete metamorphosis is that it enables the almost complete decoupling of growth and differentiation of adult structures (*8, 18*). While a trade-off between growth and differentiation is widespread in animals and plants (*18–21*), fast growth is selectively favored under many environmental conditions. These include time constraints (*22*), competition for ephemeral resources (*23*) and size or stage-specific mortality risks (*24*). In holometabolous insects, growth is confined to the larval stage and differentiation of adult organs almost entirely to the pupa, hence the growing larva should be mostly released from any constraints imposed by a trade-off between growth and differentiation (*18*). In contrast, in hemimetabolous insects, growth and differentiation occur throughout nymphal development, meaning that nymphs are almost certainly constrained by a trade-off between growth and differentiation (*20, 25*)

To investigate the hypothesis that decoupling growth and differentiation is the adaptive explanation for the evolution of complete metamorphosis, we first employ a comparative analysis to ask if holometabolous insects grow faster than hemimetabolous insects. We then implement the trade-off between growth and differentiation into a mathematical model that demonstrates how the trade-off could result in the evolution of a holometabolous lifecycle. We explore the conditions under which the transition from hemi-to holometaboly can arise.

## Comparative analysis of growth rates

For our comparative analysis, we used a database with an entomological/faunistic focus covering studies that were conducted under comparable conditions (see methods in supplement), to reduce variation caused by environmental factors. In total, we obtained data for 33 insect species, including 21 hemimetabolous and 12 holometabolous insects (figure 1, figure S1, S2). Our results show significantly higher relative growth rates (RGR) in holometabolous compared to hemimetabolous insects (RGR = 0.0548; 95% CI = 0.0109, 0.0986; p=0.016; figure 1 and tables S2, S3). Variance heterogeneity was high (I^2^ = 96.57%) and differed significantly between hemimetabolous and holometabolous insects (Breusch-Pegan test, p<0.0001; see table S4). The percentage of variation attributed to the type of metamorphosis (R^2^) was 20.79% in RGR. Half of the variance was explained by phylogeny and the other half was explained by differences among species (table S4). The maximum growth estimates of holometabolous insects were much higher than those of hemimetabolous insects (see figure 1 and table S7).

**Figure 1:**
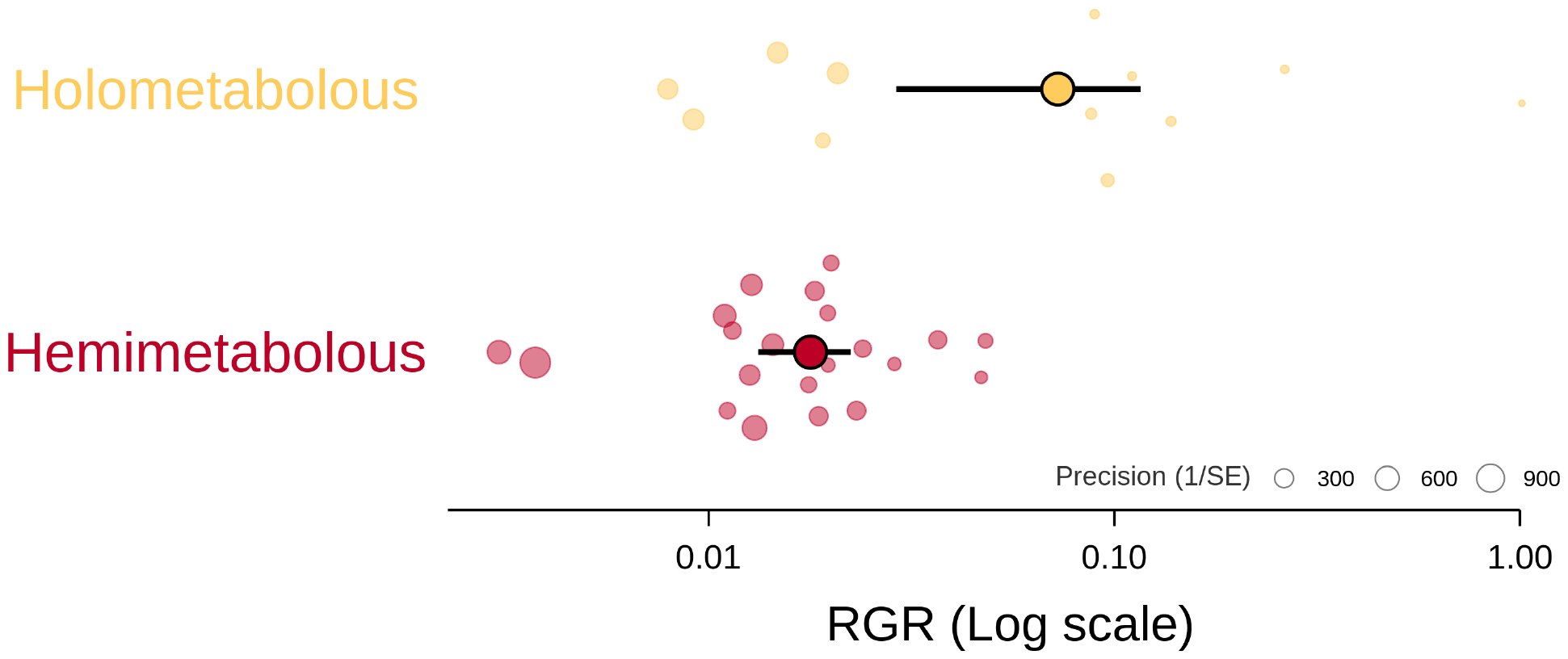
Differences in growth rates between hemi-and holometabolous insects. Mean relative growth rate (MGR, averaged over all immature moults and controlled for developmental times) per day among hemimetabolous- and holometabolous insects (RGR = 0.0548; 95% CI = 0.0109, 0.0986; p=0.016). Filled circles with black borders indicate mean growth estimates, with error bars depicting 95% confidence. Each circle without a black border represents the values of a particular insect species (scaled by their precision). Note: RGR is plotted on a log 10 scale.

## Modelling how decoupling favors growth

Given that the comparative data showed that holometabolous insects grow faster than hemimetabolous insects, we next investigated the conditions under which complete metamorphosis may have evolved. Specifically, we explored how this unique form of metamorphosis including a pupal stage might have been shaped by a trade-off between growth and differentiation. To do this, we developed a minimal model of an insect life cycle with a growth phase of duration *t* during which allocation towards differentiation can occur, followed by a (facultative) phase of pure differentiation, of duration Δ (Figure 2). Both growth and differentiation saturate, reflecting the existence of an asymptotic size, L_T_, and a final state of differentiation beyond which further differentiation is impossible.

**Figure 2:**
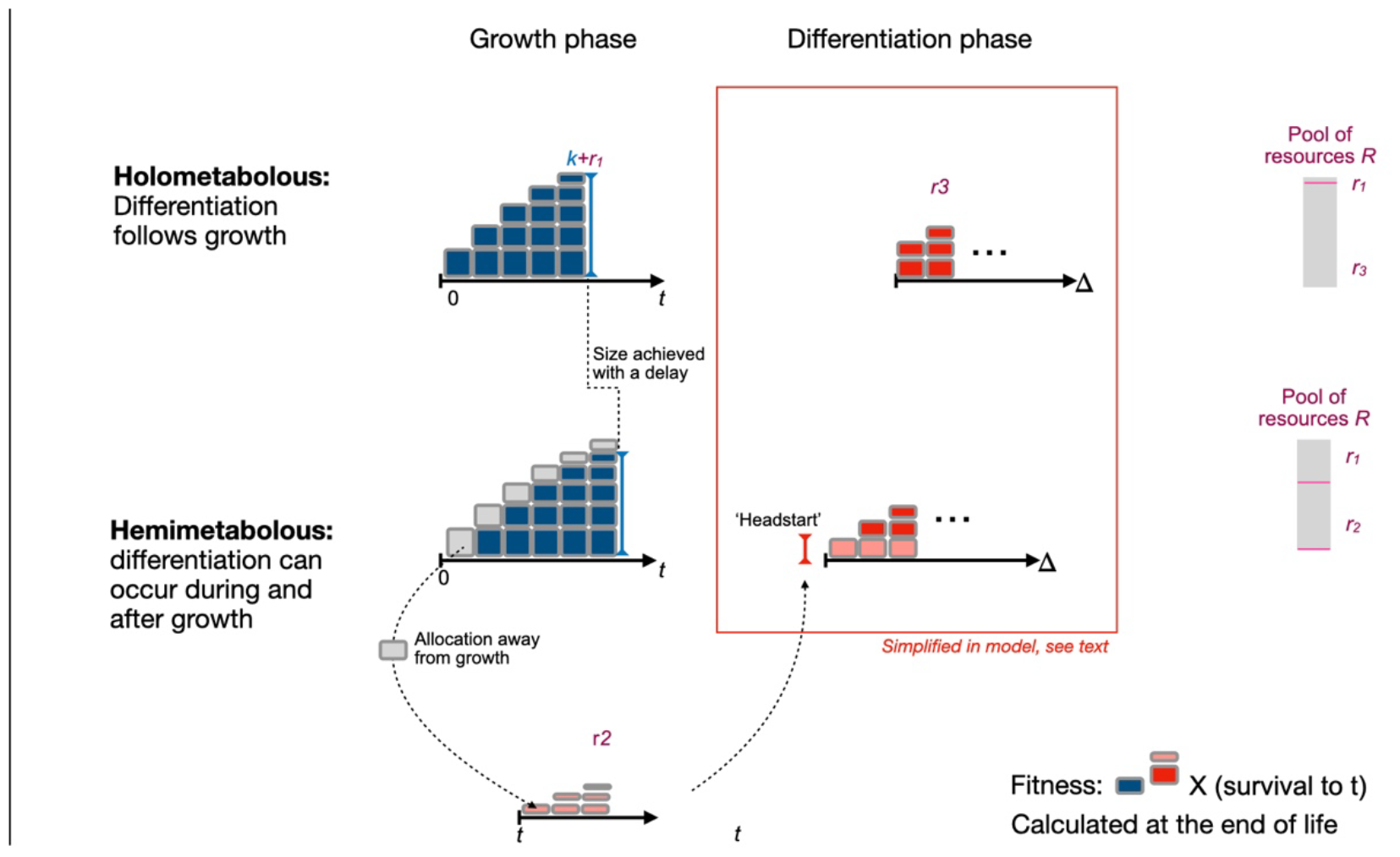
Schematic of the model. Organisms can adopt a holometabolous (top panel) or hemimetabolous (bottom panel) life cycle. In the former, a period of growth in size (accumulation of blue bricks; size at *t* is defined by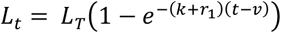, is followed by a period of differentiation (accumulation of red bricks, simplified to *r*_3_, see text). *k* describes the background growth efficiency realized in the absence of resource allocation, *r*_3_ quantifies the resource allocation towards growth in size, and *v* captures the hypothetical age at which size would be zero (required to prevent negative growth). For hemimetabolous life cycles, differentiation can also occur during the growth phase (pink bricks, lower row), by allocation of *r*_2_ resources. This reduces resources available for the rate of growth in size (greyed out bricks, upper row). During the final life phase in a hemimetabolous life cycle, remaining resources, *r*_3_, can be used to make up remaining differentiation possible, 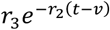, noting that for the holometabolous life cycle, *r*_2_ = 0, so this reduces to *r*_3_. Our model reflects a simplification of the full process of differentiation over time, as is captured by the inset red box. With limited data to inform this part of the model, we collapsed this down to a single phenomenon of allocation without explicitly encompassing the duration of mortality during the phase of differentiation. The total resources pool is constrained such that *r*_1_ + *r*_2_ + *r*_3_ = 1 (schematic, bottom right). Growth in size and differentiation are then combined to define fertility, and this is multiplied by survival during the growth phase to define fitness.

We found that the holometabolous life cycle (resource allocation towards differentiation absent during the growth phase*r*_2_ = 0 and present after the growth phase *r*_3_ > 0) evolves at high rates of background growth rate, *k* (Figure 3, 3^rd^ and 4^th^ panels). Increasing *k* is also associated with a declining duration of the growth phase *t* (Figure 3, 1^st^ panel) (*26*). Furthermore, the threshold value of *k* for the transition to a holometabolous life cycle declines with increasing mortality (Figure 4), as increasing mortality diminishes returns from investment in the growth phase. Qualitative results are invariant to the magnitude of *L*_*T*_ and *v*.

**Figure 3:**
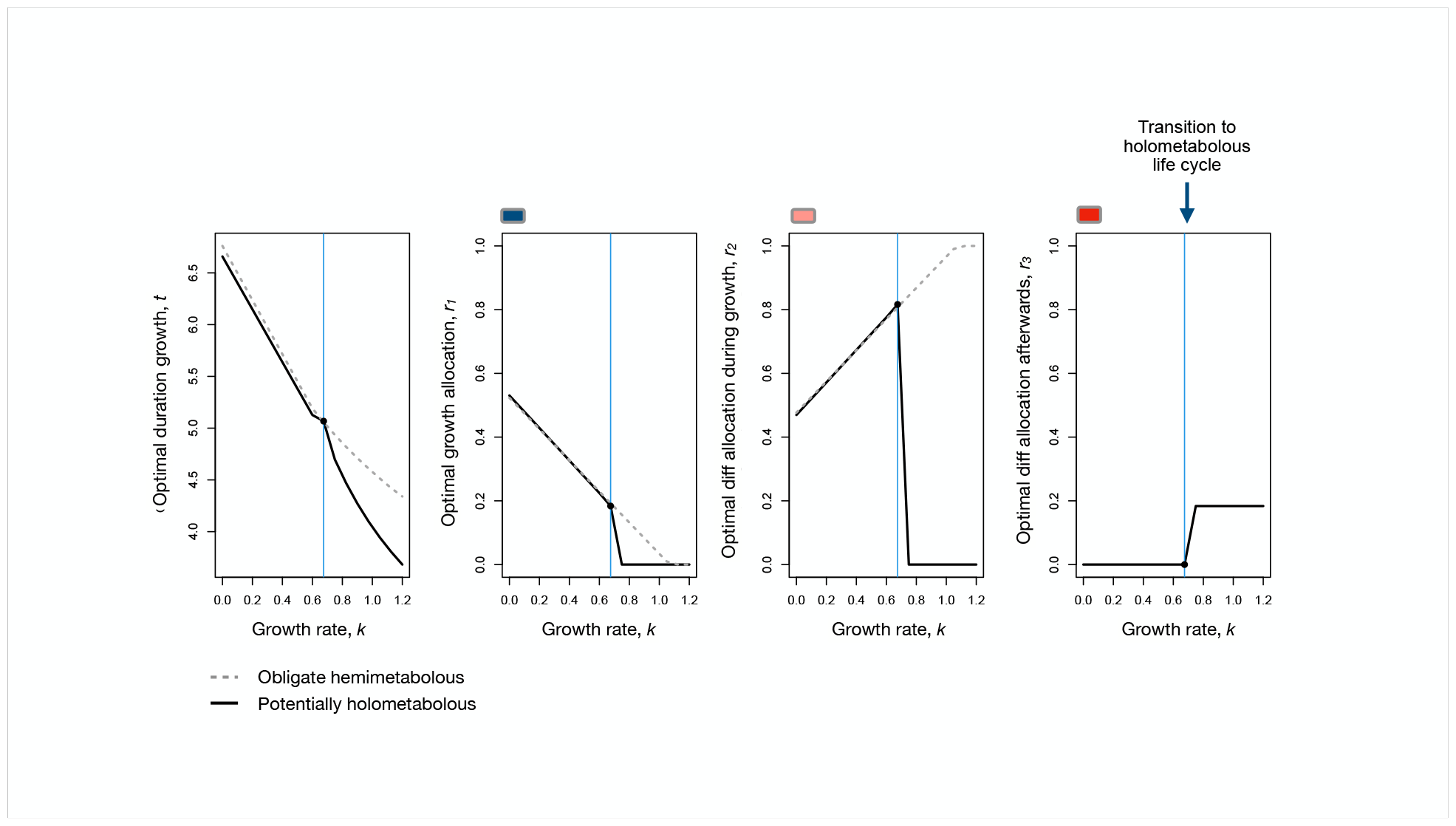
Optimal resource allocation. We numerically identified the duration of the growth phase, *t* (left panel), and allocation towards growth and differentiation during the growth phase (*r*_*1*_ and *r*_*2*_, from which *r*_*3*_ follows; center and right panels) that maximizes fitness, R_0_, across a range of values of *k*, the background growth efficiency. To identify conditions that result in the evolution of a holometabolous life cycle, we contrasted the full optimization (solid line) with optimal parameter values for an obligate hemimetabolous life cycle, i.e., where *r*_*3*_=0 (dashed lines). While background growth efficiency, *k*, increases, the optimal duration of the growth phase, *t*, declines, as individuals can reach the asymptotic size faster. Optimal allocation of resources towards the growth rate *r*_*1*_, also declines, as allocation towards differentiation during growth *r*_*2*_, increases. However, for the flexible life cycle (solid line) this quantity eventually collapses to zero as the growth rate is so high and thus the duration of the growth phase is so low, that resources for differentiation are optimally allocated entirely towards the final differentiation phase, *r*_*3*_ reflecting a transition to a holometabolous life cycle (vertical blue line). For the obligate hemimetabolous life cycle (dashed line) the patterns are monotonic along the x-axis. These results reflect a mortality rate of *μ*_*G*_ = 0. 01.

**Figure 4:**
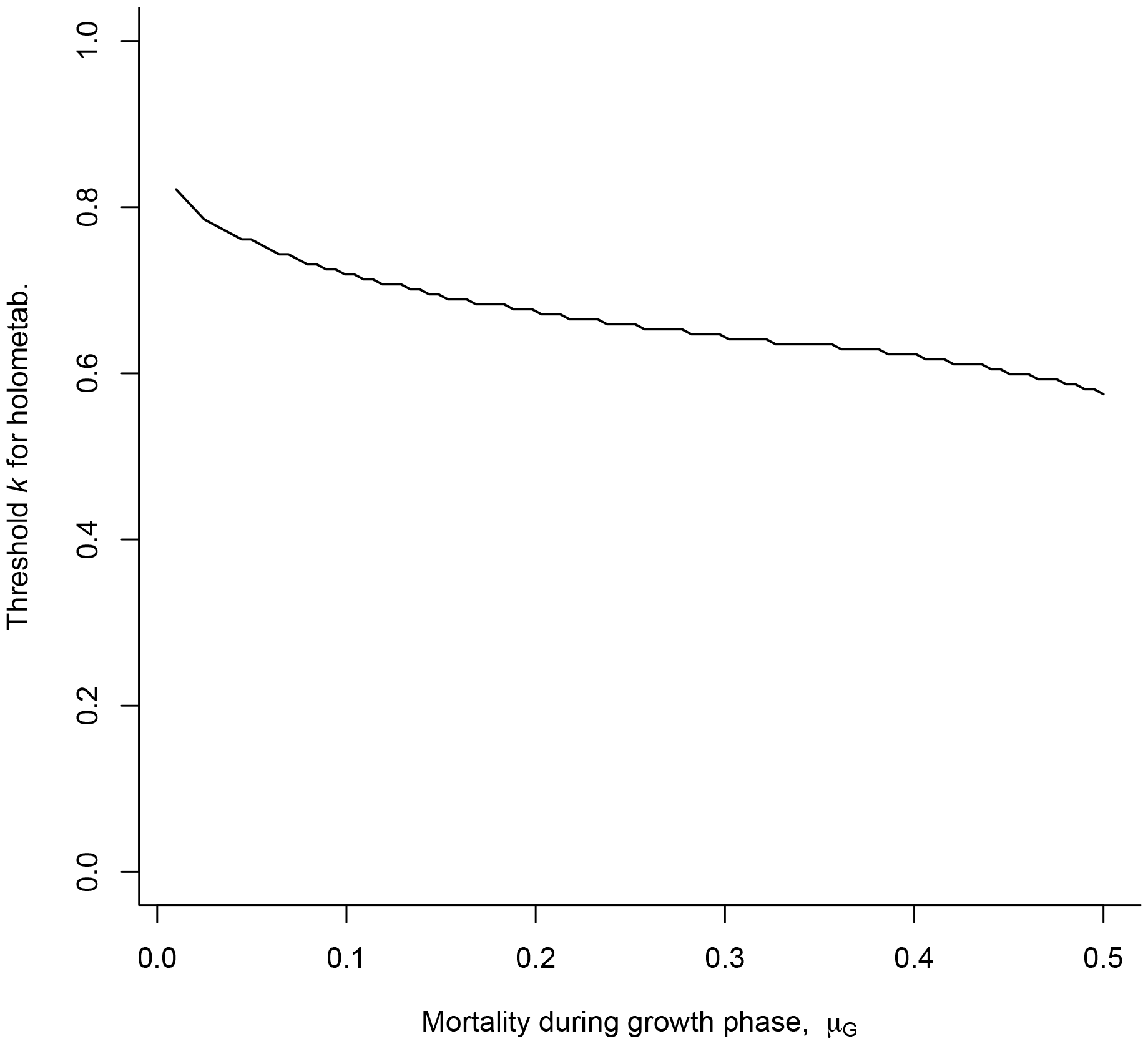
Threshold growth rate for the transition to a holometabolous life cycle under increasing mortality. As mortality during the growth phase μ_*G*_increases, the fitness contribution of the growth phase declines. As a result, the threshold growth rate *k* at which a transition to holometabolous life-cycle is expected, declines.

## Discussion

Overall, we find that holometabolous insects have higher growth rates than hemimetabolous insects. This is consistent with the fastest reported growth rates in insects in black soldier flies (*27*) and burying beetles (*28*), both of which are holometabolous. Our mathematical model shows that in the presence of a trade-off between growth and differentiation, selection for fast growth results in the temporal decoupling of growth and differentiation and can result in the evolution of holometaboly. This effect is exacerbated under increasing risk of mortality. Taken together our findings are highly consistent with the idea that holometaboly enables insects to escape the developmental constraints imposed by a trade-off between growth and differentiation.

Further support for the idea that a growth-differentiation trade-off selects for decoupling is provided by species that forego metamorphosis at least partially and obtain high growth rates. This for example applies to the neotenic Larvaceans, a group of tunicates, which have one of the fastest growth rates of all animals (*29*). Similar results have also been reported in salamanders where paedomorphic individuals, individuals that forego metamorphosis, enjoy an advantage under good growth conditions (*30*). Within the holometabolous insects a paedomorphic life style evolved independently a few times (*31*), for example in the parasitic Strepsiptera. Here females forego the pupal stage and attain a much higher mass at sexual maturity than fully differentiated males, without having a much longer development time. In all of these species, fast growth is achieved by reducing differentiation into adult forms except for the reproductive organs. As the ancestral state of insects is winged to allow for adult dispersal, simply reducing differentiation of adult structures such as the wings seems not viable in most ecological situations. The Strepsiptera mentioned above for example are endoparasites of winged insects (e.g. wasps), so dispersal for female strepsipterans is available.

Foregoing differentiation almost completely during growth in the holometabolous insects seems to have been enabled by a prolonged embryonic stage that forms the larval stage in holometabolous insects (*32*). Berlese proposed this extension of the embryonic stage in 1913 (*33*). From this statement it follows that the pupa is homologous to the nymph of hemimetabolous insects. This view is strongly supported by recent work on the main transcription factors Chinmo, Broad and E93, which together determine the developmental transitions from larva to pupa to adult (*34–37*), though alternative views exist, namely that the pupa is homologous to the last nymphal instar (*34, 38*). Our results do not make a statement about the developmental pathways that result in a larval and pupal stage of holometabolous insects, but rather identify how selection can result in decoupling growth and differentiation. Bringing together the developmental and the adaptive evolution perspective on complete metamorphosis will be an important future challenge. This should allow for understanding genetic changes and their order that were required to result in this key innovation of the pupa (*39*). It is worth noting that complete metamorphosis in the holometabolous insects happens when growth has ceased. In most other taxa, most growth takes place in the adult phenotype.

Selection for decoupling growth and differentiation does not exclude a role for existing (albeit rare (*10*)) alternate adaptive explanations for the evolution of complete metamorphosis (*8, 40, 41*). (a) Hinton proposed that the pupal stage would allow for larval stages without wing pads (*42*), which would be beneficial in burrowing insects, or also for life under bark (*43*). However, a phylogenetic reconstruction concludes that the ancestral holometabolan larva was neither burrowing nor living in crevices (*44*). (b) Wigglesworth in 1954 suggested that genetic independence between the larval and adult stages could be facilitated by the pupal stage (*45*)(*46, 47*). The literature on genetic correlations between larval and adult holometabolous insects is relatively limited, and shows varied patterns (while in fruit flies heat resistance is decoupled (*47*) between larvae and adults, antimicrobial peptide expression is not (*46*)) leaving this an open question. A final recent hypothesis is that as complete metamorphosis entails the renewal of the gut, it enables organisms to exchange their gut microbiota when they change habitat and/or diet (*40, 41*), providing an additional adaptive benefit.

How complete metamorphosis is related to the evolutionary success of the holometabolous insects, measured in terms of both species richness and habitat dominance, is poorly understood (*8, 48*). One explanation is that once complete metamorphosis had evolved, the resulting higher modularity would enable higher evolvability, as for example shown by the high diversity of mouthparts in holometabolous insects (*49*). If, as we propose here, the decoupling of growth and differentiation has been the main driver for the evolution of complete metamorphosis, then any ecological situation where fast growth is beneficial would give holometabolous insects a significant competitive advantage over other organisms that display alternative forms of metamorphosis or no metamorphosis at all. The breaking of constraints on growth almost certainly provides an added evolutionary benefit to the other potential advantages provided by biphasic metamorphic life cycles.

## Acknowledgments

We are grateful to Oren Harman and Dino McMahon for comments on the manuscript.

## Funding

DFG, FOR 5026 (JR)

DFG RO 2284/4-1 (CM, JR)

FU Berlin (CJEM)

DFG Heisenberg 430170797 (UKS)

## Data and materials availability

All data and code are available in the main text or the supplementary materials.

## References and Notes

1. N. A. Moran, Adaptation and Constraint in the Complex Life Cycles of Animals. Annual Review of Ecology and Systematics 2, 573–600 (1994).

2. E. E. Werner, Amphibian Metamorphosis: Growth Rate, Predation Risk, and the Optimal Size at Transformation. The American Naturalist 128, 319–341 (1986).

3. H. ten Brink, A. M. de Roos, U. Dieckmann, The Evolutionary Ecology of Metamorphosis. The American Naturalist 193, E116–E131 (2019).

4. C. Mora, D. P. Tittensor, S. Adl, A. G. B. Simpson, B. Worm, How Many Species Are There on Earth and in the Ocean? PLoS Biol 9, e1001127 (2011).

5. B. Misof, S. Liu, K. Meusemann, R. S. Peters, A. Donath, C. Mayer, P. B. Frandsen, J. Ware, T. Flouri, R. G. Beutel, O. Niehuis, M. Petersen, F. Izquierdo-Carrasco, T. Wappler, J. Rust, A. J. Aberer, U. Aspöck, H. Aspöck, D. Bartel, A. Blanke, S. Berger, A. Böhm, T. R. Buckley, B. Calcott, J. Chen, F. Friedrich, M. Fukui, M. Fujita, C. Greve, P. Grobe, S. Gu, Y. Huang, L. S. Jermiin, A. Y. Kawahara, L. Krogmann, M. Kubiak, R. Lanfear, H. Letsch, Y. Li, Z. Li, J. Li, H. Lu, R. Machida, Y. Mashimo, P. Kapli, D. D. McKenna, G. Meng, Y. Nakagaki, J. L. Navarrete-Heredia, M. Ott, Y. Ou, G. Pass, L. Podsiadlowski, H. Pohl, B. M. Von Reumont, K. Schütte, K. Sekiya, S. Shimizu, A. Slipinski, A. Stamatakis, W. Song, X. Su, N. U. Szucsich, M. Tan, X. Tan, M. Tang, J. Tang, G. Timelthaler, S. Tomizuka, M. Trautwein, X. Tong, T. Uchifune, M. G. Walzl, B. M. Wiegmann, J. Wilbrandt, B. Wipfler, T. K. F. Wong, Q. Wu, G. Wu, Y. Xie, S. Yang, Q. Yang, D. K. Yeates, K. Yoshizawa, Q. Zhang, R. Zhang, W. Zhang, Y. Zhang, J. Zhao, C. Zhou, L. Zhou, T. Ziesmann, S. Zou, Y. Li, X. Xu, Y. Zhang, H. Yang, J. Wang, J. Wang, K. M. Kjer, X. Zhou, Phylogenomics resolves the timing and pattern of insect evolution. Science 346, 763–767 (2014).

6. B. Wang, C. Xu, E. A. Jarzembowski, Ecological radiations of insects in the Mesozoic. Trends in Ecology & Evolution, S0169534722000441 (2022).

7. D. A. Grimaldi, M. Engel, The Evolution of the Insects (Cambridge University Press, Cambridge, 2005).

8. J. Rolff, P. R. Johnston, S. Reynolds, Complete metamorphosis of insects. Phil. Trans. R. Soc. B 374, 20190063 (2019).

9. M. J. R. Hall, D. Martín-Vega, Visualization of insect metamorphosis. Phil. Trans. R. Soc. B 374, 20190071 (2019).

10. X. Bellés, Insect Metamorphosis: From Natural History to Regulation of Development and Evolution (Academic Press, London, 2020).

11. S. Aldaz, L. Escudero, Imaginal discs. Current Biology, R429–R431.

12. R. Chapman, The Insects - Structure and Function (Cambridge University Press, Cambridge, 2013).

13. F. Sehnal, Morphology of Insect Development.

14. H. E. Hinton, THE ORIGIN AND FUNCTION OF THE PUPAL STAGE. Proceedings of the Royal Entomological Society of London. Series A, General Entomology 38, 77–85 (1963).

15. J. W. Truman, L. M. Riddiford, The evolution of insect metamorphosis: a developmental and endocrine view. Phil. Trans. R. Soc. B 374, 20190070 (2019).

16. S. Reynolds, Cooking up the perfect insect: Aristotle’s transformational idea about the complete metamorphosis of insects. Phil. Trans. R. Soc. B 374, 20190074 (2019).

17. M. A. Albecker, L. G. E. Wilkins, S. A. Krueger-Hadfield, S. M. Bashevkin, M. W. Hahn, M. P. Hare, H. K. Kindsvater, M. A. Sewell, K. E. Lotterhos, A. M. Reitzel, Does a complex life cycle affect adaptation to environmental change? Genome-informed insights for characterizing selection across complex life cycle. Proc. R. Soc. B. 288, 20212122 (2021).

18. J. D. Arendt, Adaptive Intrinsic Growth Rates: An Integration Across Taxa. The Quarterly Review of Biology 72, 149–177 (1997).

19. D. A. Herms, W. J. Mattson, The Dilemma of Plants: To Grow or Defend. The Quarterly Review of Biology 67, 283–335 (1992).

20. J. Rolff, F. V. de Meutter, R. Stoks, Time Constraints Decouple Age and Size at Maturity and Physiological Traits. 7.

21. C. M. Dmitriew, The evolution of growth trajectories: what limits growth rate? Biological Reviews 86, 97–116 (2011).

22. L. Rowe, D. Ludwig, Size and timing of metamorphosis in complex life cycles: times constraints and variation. Ecology, 413–427.

23. L. D. Mueller, Density-Dependent Population Growth and Natural Selection in Food-Limited Environments: The Drosophila Model. The American Naturalist 132, 786–809 (1988).

24. K. Spitze, CHAOBORUS PREDATION AND LIFE‐HISTORY EVOLUTION IN DAPHNIA PULEX: TEMPORAL PATTERN OF POPULATION DIVERSITY, FITNESS, AND MEAN LIFE HISTORY. Evolution 45, 82–92 (1991).

25. P. T. Rühr, T. van de Kamp, T. Faragó, J. U. Hammel, F. Wilde, E. Borisova, C. Edel, M. Frenzel, T. Baumbach, A. Blanke, Juvenile ecology drives adult morphology in two insect orders. Proc. R. Soc. B. 288, 20210616 (2021).

26. C. J. E. Metcalf, B. Koskella, Protective microbiomes can limit the evolution of host pathogen defense. Evolution Letters 3, 534–543 (2019).

27. I. J. Banks, W. T. Gibson, M. M. Cameron, Growth rates of black soldier fly larvae fed on fresh human faeces and their implication for improving sanitation. Tropical Med Int Health 19, 14–22 (2014).

28. P. T. Smiseth, C. T. Darwell, A. J. Moore, Partial begging: an empirical model for the early evolution of offspring signalling. Proc. R. Soc. Lond. B 270, 1773–1777 (2003).

29. C. Jaspers, R. R. Hopcroft, T. Kiørboe, F. Lombard, Á. López-Urrutia, J. D. Everett, A. J. Richardson, Gelatinous larvacean zooplankton can enhance trophic transfer and carbon sequestration. Trends in Ecology & Evolution 38, 980–993 (2023).

30. M. Denoël, P. Joly, H. H. Whiteman, Evolutionary ecology of facultative paedomorphosis in newts and salamanders. Biological Reviews 80, 663–671 (2005).

31. D. P. Mcmahon, A. Hayward, Why grow up? A perspective on insect strategies to avoid metamorphosis. Ecological Entomology 41, 505–515 (2016).

32. J. W. Truman, L. M. Riddiford, Endocrine Insights into the Evolution of Metamorphosis in Insects. Annu. Rev. Entomol. 47, 467–500 (2002).

33. A. Berlese, Intorno alle metamorfosi degli insetti. Redia 9, 121–136. 119, e2204972119 (2022).

34. S. E. Reynolds, A transcription factor that enables metamorphosis. Proc. Natl. Acad. Sci. U.S.A. 119, e2204972119 (2022).

35. J. W. Truman, L. M. Riddiford, Chinmo is the larval member of the molecular trinity that directs Drosophila metamorphosis. Proc. Natl. Acad. Sci. U.S.A. 119, e2201071119 (2022).

36. A. Fernandez-Nicolas, G. Machaj, A. Ventos-Alfonso, V. Pagone, T. Minemura, T. Ohde, T. Daimon, G. Ylla, X. Belles, Reduction of embryonic E93 expression as a hypothetical driver of the evolution of insect metamorphosis. Proc. Natl. Acad. Sci. U.S.A. 120, e2216640120 (2023).

37. J. W. Truman, The Evolution of Insect Metamorphosis. Current Biology 29, R1252–R1268 (2019).

38. M. Jindra, Where did the pupa come from? The timing of juvenile hormone signalling supports homology between stages of hemimetabolous and holometabolous insects. Phil. Trans. R. Soc. B 374, 20190064 (2019).

39. S. Stankowski, Z. B. Zagrodzka, M. D. Garlovsky, A. Pal, D. Shipilina, D. G. Castillo, H. Lifchitz, A. L. Moan, E. Leder, J. Reeve, K. Johannesson, A. M. Westram, R. K. Butlin, The genetic basis of a recent transition to live-bearing in marine snails. (2024).

40. C. Manthey, P. R. Johnston, S. Nakagawa, J. Rolff, Complete metamorphosis and microbiota turnover in insects. Molecular Ecology, mec.16673 (2022).

41. T. J. Hammer, N. A. Moran, Links between metamorphosis and symbiosis in holometabolous insects. Phil. Trans. R. Soc. B 374, 20190068 (2019).

42. H. Hinton, On the origin and function of the pupal stage. Transactions of the Royal Entomological Society 99, 395–409.

43. W. D. Hamilton, “Funeral Feasts: Evolution and diversity under barke” In Narrow Roads of Gene Land (WH Freeman, New York, NY, ed. 1, 1996)vol. 1, pp. 387–422.

44. R. S. Peters, K. Meusemann, M. Petersen, C. Mayer, J. Wilbrandt, T. Ziesmann, A. Donath, K. M. Kjer, U. Aspöck, H. Aspöck, A. Aberer, A. Stamatakis, F. Friedrich, F. Hünefeld, O. Niehuis, R. G. Beutel, B. Misof, The evolutionary history of holometabolous insects inferred from transcriptome-based phylogeny and comprehensive morphological data. BMC Evol Biol 14, 52 (2014).

45. V. Wigglesworth, The Physiology of Insect Metamorphosis (Cambridge University Press, Cambridge, 1954)vol. 1 of Cambridge Monographs in experimental Biology I.

46. S. Fellous, B. P. Lazzaro, Potential for evolutionary coupling and decoupling of larval and adult immune gene expression. Molecular Ecology 20, 1558–1567 (2011).

47. V. Loeschcke, R. A. Krebs, SELECTION FOR HEAT-SHOCK RESISTANCE IN LARVAL AND IN ADULT DROSOPHILA BUZZATII: COMPARING DIRECT AND INDIRECT RESPONSES. Evolution 50, 2354–2359 (1996).

48. D. B. Nicholson, A. J. Ross, P. J. Mayhew, Fossil evidence for key innovations in the evolution of insect diversity. Proc. R. Soc. B. 281, 20141823 (2014).

49. A. S. Yang, Modularity, evolvability, and adaptive radiations: a comparison of the hemi‐ and holometabolous insects. Evolution and Development 3, 59–72 (2001).

