## Supplementary material for "Rapid growth and the evolution of complete metamorphosis in insects": SupplementMantheyetal.docx

Supplement

Materials and Methods

Supplementary Text

Figs. S1 to S2

Tables S1 to S9

References (50–57)

R-scripts

Movies NA

Audio NA

Data NA

**Supplement**

**Methods**

**Literature Search**

Publications were collected that reported total body lengths and developmental times for immature stages of insects. Literature was systematically searched on ZOBODAT, the Zoological-Botanical Database (www.zobodat.at), which contains digitised literature on insect studies, including studies published in German. Titles were searched exclusively. The search terms included all 29 order names of insects (Misof et al., 2014), their respective German names and corresponding synonyms (see supplementary table 1). Publications with the following words or phrases in their titles were not further reviewed: Aufzählung (enumeration/list); Verzeichnis (index); Classification; neue Arten (new species)/ neue exotische (new exotic) / neue Formen (new forms); Bemerkungen über (remarks about) / Bemerkungen zur (remarks on); Beiträge zur Kenntnis (contributions to knowledge); Bestimmungsschüssel (identification key); Ergebnisse zoologischer Reise (results of zoological trip); Forschungsreise (research trip); Beschreibung einiger (description of some); Zur Verbreitung von (on the distribution of). The remaining publications were further investigated. Publications were included in the final dataset if they met all the following criteria:

(i) Living animals or exuviae were measured

(ii) Total body lengths from all or most immature stages were taken

(iii)Developmental times from all collected immature stages were reported.

Publications dates ranged from 1904 to 2015. The majority of studies from the literature were conducted under comparable conditions to reduce variation caused by environmental factors such as climate (table 8)."Table 1 in the supplement lists the number of generated publications from the initial and refined search and the species numbers for each insect order (see also figure 1).

**Growth rates - Empirical data**

Mean relative growth rates (RGR) per insect species were estimated. RGR is a standardised measure of growth in insects (e.g. Tammaru et al. 2004) which is the ratio of the size of one instar to the size of the previous instar relative to size, taking into account the differences in initial sizes and can, therefore, be used to compare the growth of different species. The growth ratios for each species were averaged over all immature moults and controlled for developmental times. The RGR was calculated as follows:

$$RGR=\frac{1}{n}\sum_{i=1}^{n} (\frac{ln({\text{size instar}_{n}}/{\text{size instar}_{n-1}})}{\Delta t})=(\frac{ln(\text{size instar}_{n})-ln(\text{size instar}_{n-1})}{\Delta t})$$

Where n is the number of instars for a species of concern, and t is the developmental time - the instar duration in days. Non-feeding stages where larvae entered a wandering stage were excluded. Total body lengths were used as size measures. The peak larval length was taken as the final instar size. The sizes of three Odonates (Erythemis simplicicollis, Anax junius and Nasiaeshna pentacantha) based on exuviae length measures. Weighted RGR were calculated for one hemimetabolous (Eucorydia yasumatsui) and three holometabolous species (Tenebrio molitor, Zophobas atratus and Lasiocampa pini), because of their variation in the number of instars (see supplementary table 4 for more details). The sizes of the three Gryllidae species were measured every ten days, and consequently, the RGR was calculated in ten-day intervals.

**Statistical Analysis**

All statistical analyses were performed in R (version 3.6.3; R Core Team, 2020). Differences in the RGR comparing holo- and hemimetabolous insects was tested. The variance heterogeneity (Dutilleul & Legendre, 1993) across the two groups was tested with the Breusch- Pegan test using the ncvTest function from the R package car (Fox, Weisberg, & Price, 2021). Regression analyses controlled for phylogeny were performed to reduce type I error rates. Grafen’s branch lengths were generated before modelling the phylogenetic correlation matrix for the models (Grafen, 1989). The models were phylogenetic linear mixed-effects models (Manthey et al. 2022) using the rma.mv function from the R package metafor (Cinar, Nakagawa, & Viechtbauer, 2021) that incorporates sampling variance (the square of SE). First, an intercept model with two random effects (species ID and phylogeny) was fitted. The percentage of total variation (I2) due to heterogeneity rather than chance (sampling variance) was calculated (Higgins, 2003; Senior et al., 2016) and separated into the heterogeneity explained by phylogeny and differences between observation points (species ID) using the i2_ml function from the R package orchaRd (Nakagawa et al., 2020). A second model with the two random effects (species ID and phylogeny) and the type of metamorphosis as a fixed effect that looked at the contrast between hemi- and holometabolous insects was fitted. The percentage of variation in the growth estimates attributed to the type of metamorphosis was calculated as marginal R2 (Nakagawa & Schielzeth, 2013) using the r2_ml function from the orchaRd package (Nakagawa et al., 2020). A third model that specified the variance structure of the two insect groups by modelling heteroscedasticity (heterogeneity of variances) was fitted and used to visualise the results using the orchaRd package (Nakagawa et al., 2020). The orchard plots display the group means with 95% confidence intervals, the estimates for each species scaled by their precision (1/standard error, SE) and 95% prediction intervals of the group means for hemi-and holometabolous insects.

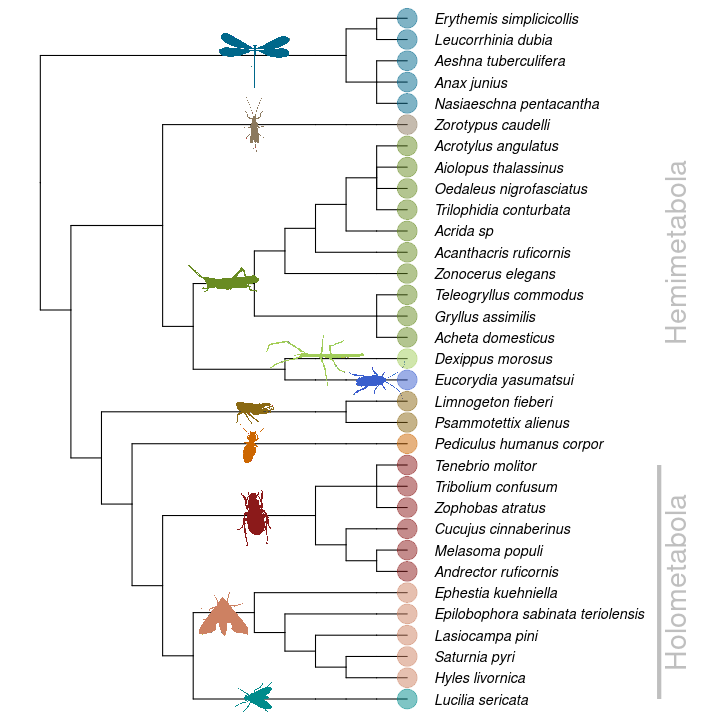
**Figure S1**: **Phylogeny of the insect species included in our comparative study of growth rates.** Our analysis includes 21 hemimetabolous and 12 holometabolous insects. The Hemimetabola cover seven orders (Odonata, Zoraptera, Orthoptera, Phasmatodea, Blattodea, Hemiptera and Psocodea) and the Holometabola three (Coleoptera, Lepidoptera, and Diptera). The growth trajectories for each species can be found in the supplementary material.

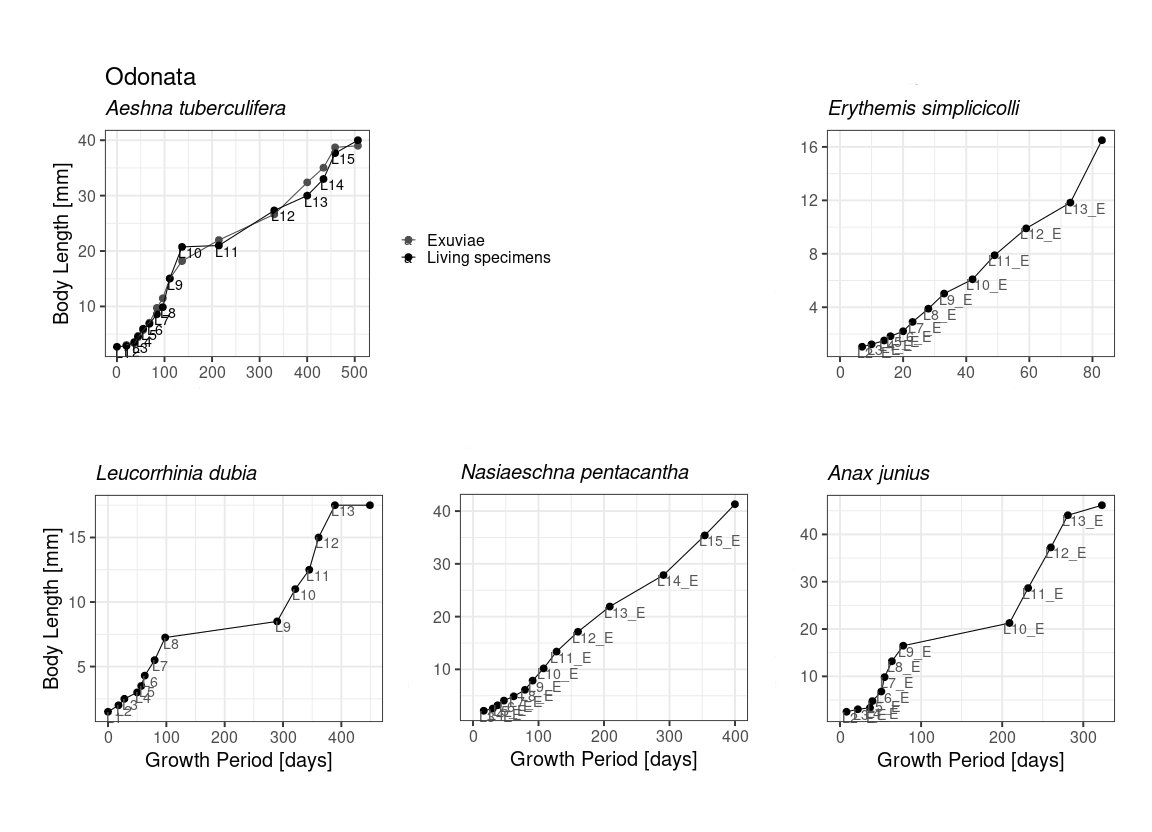

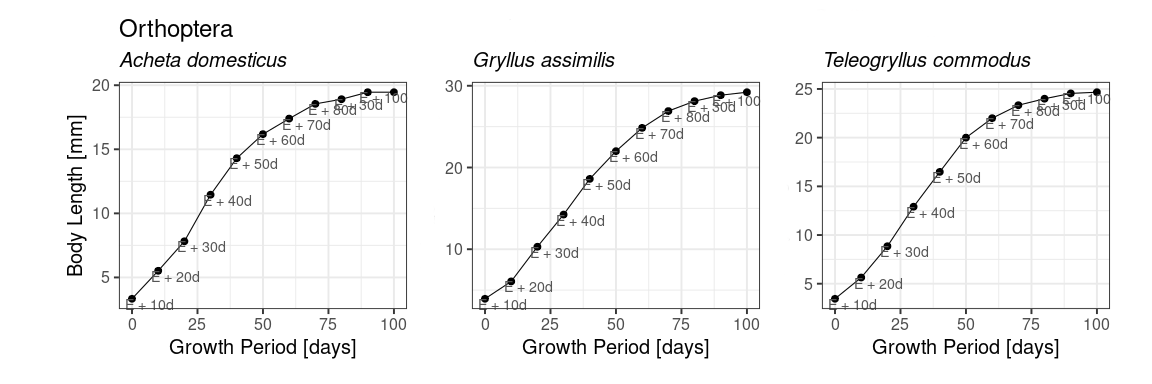

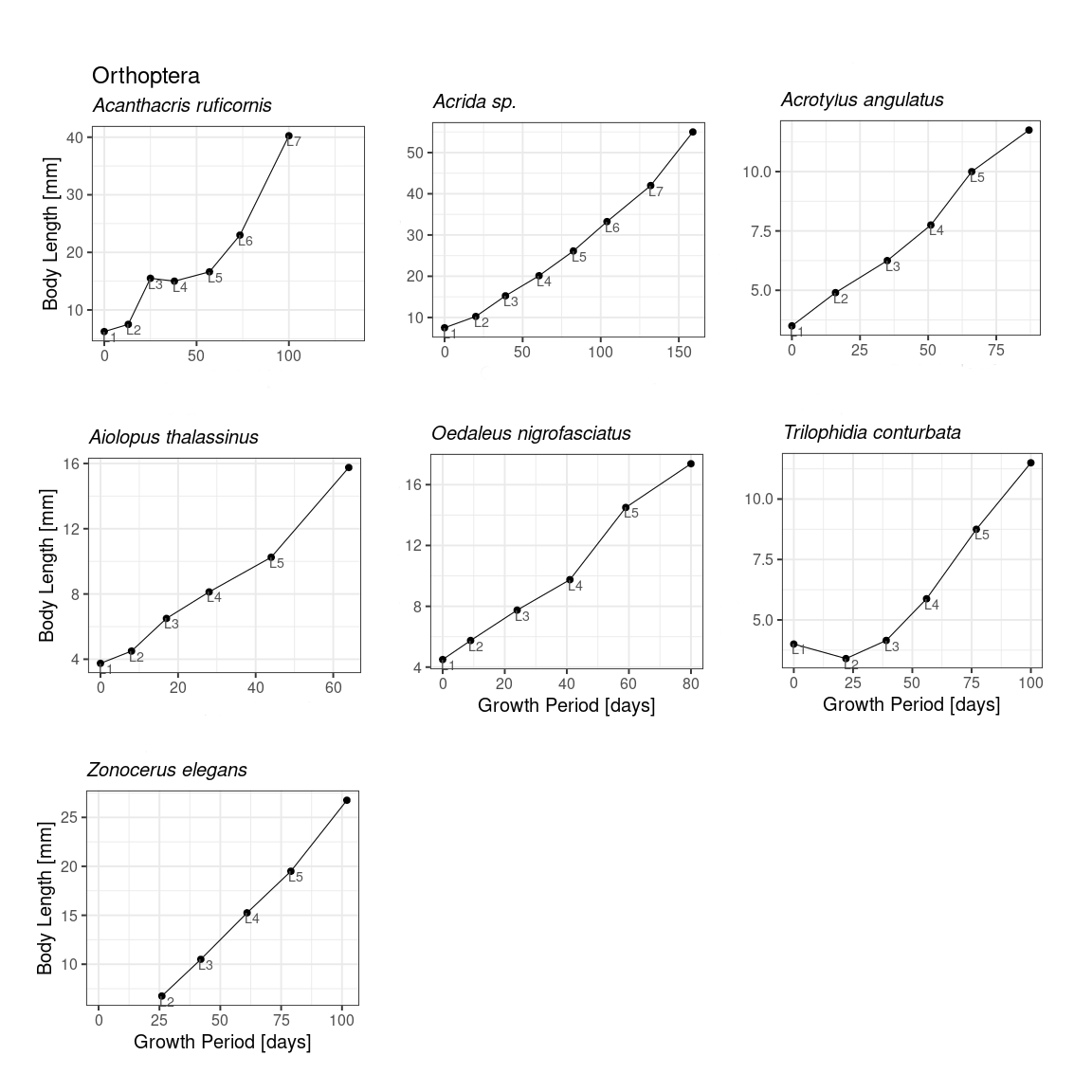

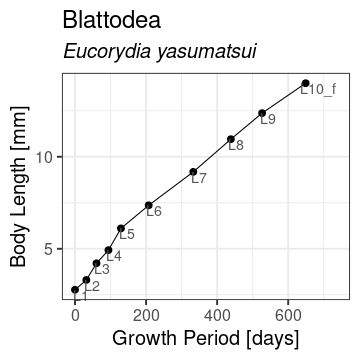

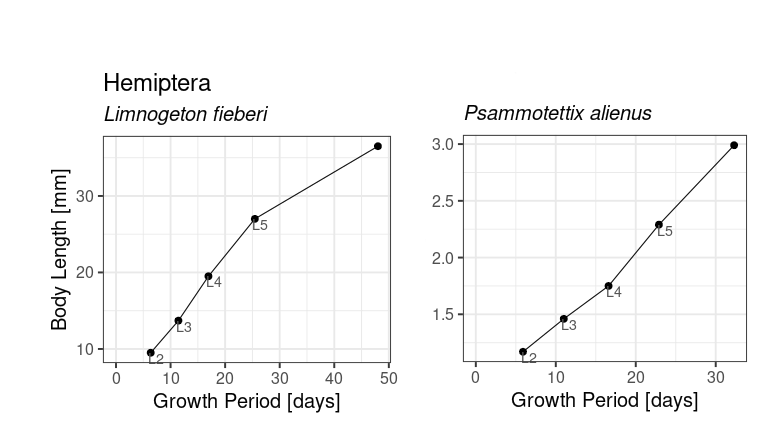

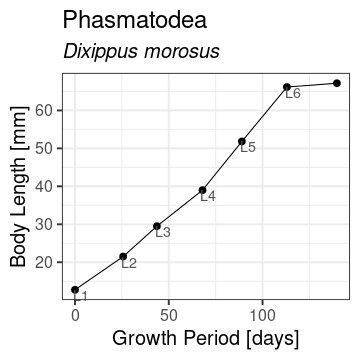

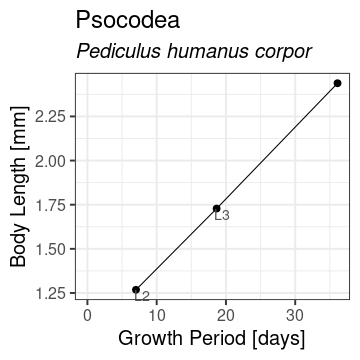

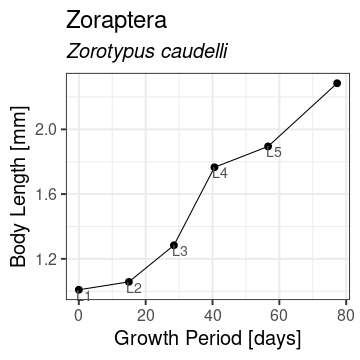

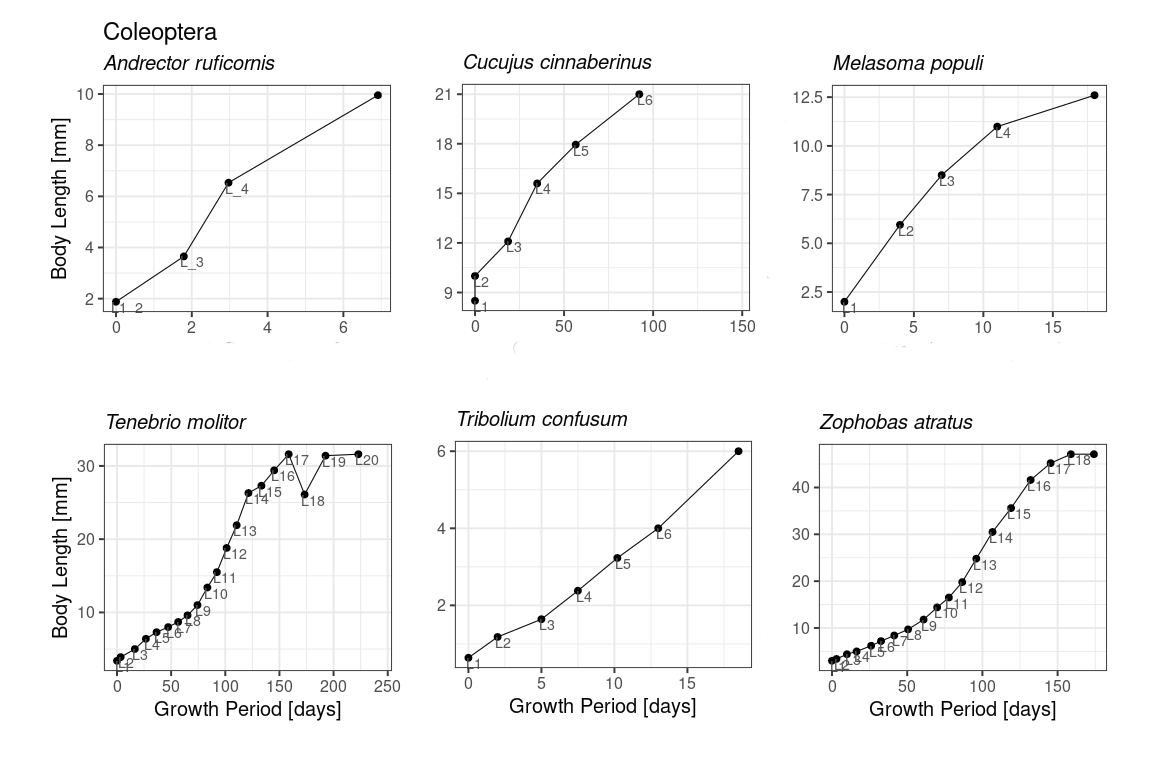

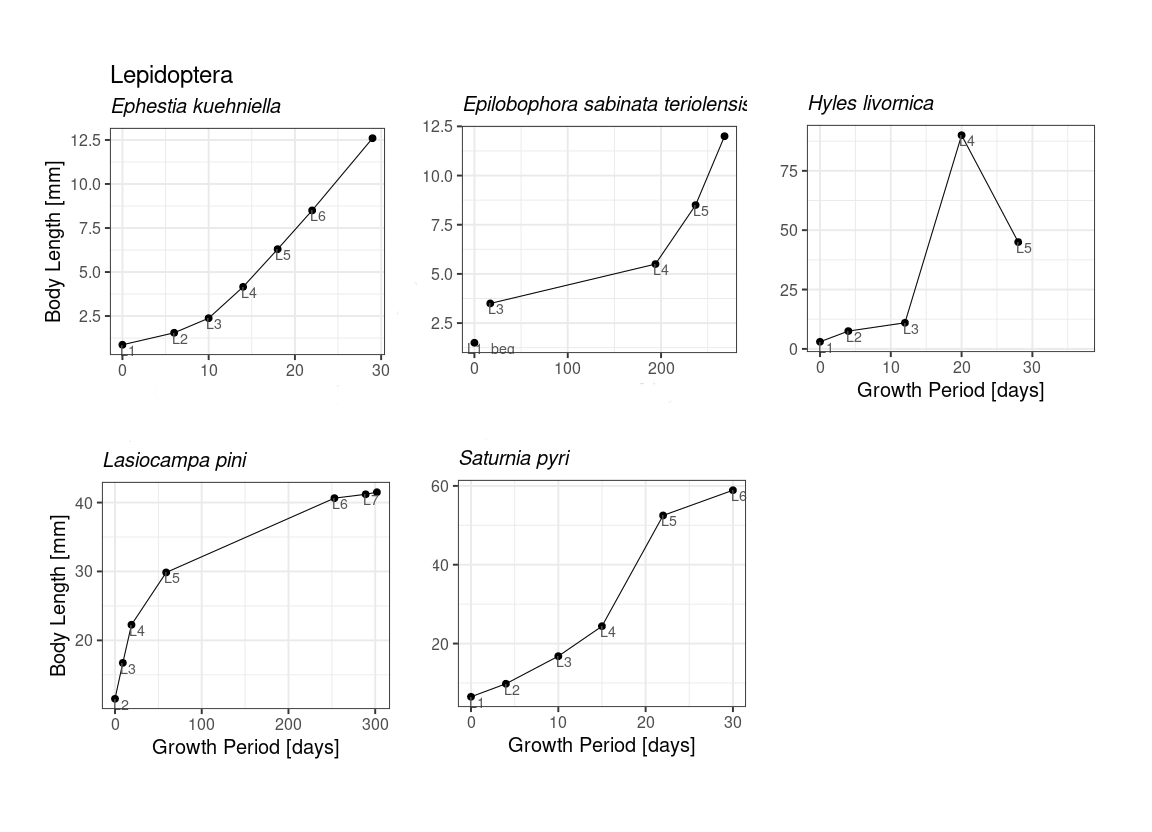

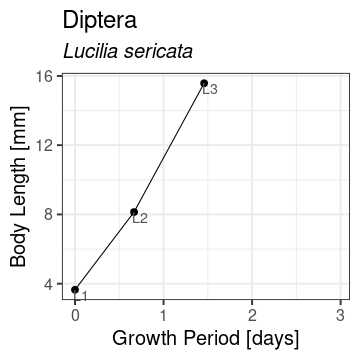

**Figure S2**: The growth trajectories of all insect species, ordered by insect order, included in the comparative analysis (mean relative growth rates per immature stage).

**Table** **S1**: Keywords used to search ZOBODAT for literature on insect growth. Also shown are the number of publications of the initial and final search (all criteria met) and the number of insects we used for the final dataset.

|  | Search terms | | No. of publications (initial search) | No. of publication that fit all criteria | Species number |
| --- | --- | --- | --- | --- | --- |
|  | Keywords (from Misof et al, 2014) | German names and synonyms |  |  |  |
| Hemimetabola | Archaeognatha | Felsenspringer | 18 | 0 | 0 |
|  | Zygentoma | Fischchen | 10 | 0 | 0 |
|  | Odonata | Libelle; Wasserjungfer | 928 | 5 | 5 |
|  | Ephemeoptera | Eintagsfliege | 80 | 0 | 0 |
|  | Zoraptera | Bodenlaus | 4 | 1 | 1 |
|  | Dermaptera | Ohrwurm | 50 | 0 | 0 |
|  | Plecoptera | Steinfliege | 546 | 0 | 0 |
|  | Orthoptera | Heuschrecke | 538 | 2 | 10 |
|  | Mantophasmatodea | Gladiatorschrecke; Gladiator; Fersenläufer | 4 | 0 | 0 |
|  | Grylloblattodea | Grillenschabe; Notoptera | 0 | 0 | 0 |
|  | Embioptera | Tarsenspinner; Fersenspinner | 4 | 0 | 0 |
|  | Phasmatodea | Gespenstschrecke; Phasmida; Phasmiden | 37 | 1 | 1 |
|  | Mantodea | Fangschrecke; Gottesanbeterin | 104 | 0 | 0 |
|  | Blattodea | Schabe; Blatteria | 61 | 1 | 1 |
|  | Isoptera | Termiten | 143 | 0 | 0 |
|  | Thysanoptera | Fransenflügler; Thrips; Blasenfüße; Gewittertierchen; Gewitterwürmer | 76 | 0 | 0 |
|  | Hemiptera | Schnabelkerfe; Rhynchota | 584 | 2 | 2 |
|  | Psocodea | Staublaus; Psocoptera | 7 | 1 | 1 |
| Holometabola | Hymenoptera | Hautflügler | 3526 | 0 | 0 |
|  | Raphidioptera | Kamelhalsfliege | 112 | 0 | 0 |
|  | Megaloptera | Großflügler; Schlammfliege | 40 | 0 | 0 |
|  | Neurotera | Netzflügler; Planipennia | 292 | 0 | 0 |
|  | Strepsiptera | Fächeflügler | 41 | 0 | 0 |
|  | Coleoptera | Käfer | 5478 | 6 | 6 |
|  | Trichoptera | Köcherfliege | 678 | 0 | 0 |
|  | Lepidopera | Schmetterling | 4791 | 5 | 5 |
|  | Siphonaptera | Flöhe | 68 | 0 | 0 |
|  | Mecoptera | Schnabelfliege; Schnabelhafte | 5 | 0 | 0 |
|  | Diptera | Zweiflügler | 2014 | 1 | 1 |

**Table** **S2**: Results of the regression analyses controlled for phylogeny (multivariate linear mixed-effects models) testing differences in *RGR* between hemi- and holometabolous insects.

| Model |  | Estimate | SE | T-value | Df | P-value | Lower CI | Upper CI |  |
| --- | --- | --- | --- | --- | --- | --- | --- | --- | --- |
| (1) Intercept model | Intercept | 0.023 | 0.003 | 7.156 | 32 | <.0001 | 0.016 | 0.029 | *** |
| (2) Model looking at contrast  between the groups | Intercept | 0.019 | 0.004 | 4.737 | 31 | <.0001 | 0.011 | 0.027 | *** |
|  | Hemi Holo | 0.018 | 0.008 | 2.286 | 31 | 0.0293 | 0.002 | 0.035 | * |
| (3) Modelling heteroscadasticity | Intercept | 0.0178 | 0.0023 | 7.9218 | 31 | <.0001 | 0.0132 | 0.0224 | *** |
|  | Hemi_Holo | 0.0548 | 0.0215 | 2.5482 | 31 | 0.0160 | 0.0109 | 0.0986 | * |

**Table** **S3**: The regression analysis results controlled for phylogeny testing differences in RGR between hemi- and holometabolous insects for the reduced dataset without the three Gryllidae species measured in ten-day intervals (multivariate linear mixed-effects models with specified variance structure by modelling heteroscedasticity).

| Growth estimate |  | Estimate | SE | T-value | Df | P-value | Lower CI | Upper CI |  |
| --- | --- | --- | --- | --- | --- | --- | --- | --- | --- |
| (3) Modelling heteroscadasticity | Intercept | 0.0179 | 0.0026 | 6.8023 | 28 | <.0001 | 0.0125 | 0.0233 | *** |
|  | Hemi_Holo | 0.0547 | 0.0215 | 2.5414 | 28 | 0.0169 | 0.0106 | 0.0988 | * |

**Table** **S4**: Results of the Breusch-Pagan test for heteroscedasticity (heterogeneity of variances) using the *ncvTest* function from the R package car and the percentage of total variation due to heterogeneity from the phylogenetic intercept models using the *i2_ml function* from the orchaRd package.

| Growth estimate | Breusch-Pegan test | | | |  | Extent of heterogeneity ( I^2^ ) | | |
| --- | --- | --- | --- | --- | --- | --- | --- | --- |
|  | X^2^ | Df | P-value |  |  | Total I^2^ [ % ] | I^2^ due to phylogeny  [ % ] | I^2^ due to species ID [ % ] |
| RGR | 28.572 | 1 | 9.0266e-08 | *** |  | 96.57 | 48.28 | 48.28 |

**Table** **S5**: Table of results for the visualised models showing the plotted group estimates with their respective confidence intervals (CI) and prediction intervals (PR).

| Visualised model | Growth | Estimate | Lower CI | Upper CI | Lower PR | Upper PR |
| --- | --- | --- | --- | --- | --- | --- |
| Heteroscadasticity model  for RGR | Hemimetabola | 0.018 | 0.013 | 0.022 | -0.001 | 0.037 |
|  | Holometabola | 0.073 | 0.029 | 0.116 | -0.067 | 0.213 |

**Table** **S6**: Listed are the insect species with a variable number of instars. Their pupation (Holometabola) and adult eclosion (Hemimetabola) rates were used to calculate weighted RGR.

|  | Insect | No. of instars | Rate [ % ] |
| --- | --- | --- | --- |
| Hemi. | *Eucorydia yasumatsui* | 8 | 27.27 |
|  |  | 9 | 68.18 |
|  |  | 10 | 4.55 |
| Holometabola | *Tenebrio molitor* | 14 | 5.63 |
|  |  | 15 | 19.39 |
|  |  | 16 | 21.98 |
|  |  | 17 | 28.32 |
|  |  | 18 | 15.05 |
|  |  | 19 | 6.29 |
|  |  | 20 | 2.5 |
|  | *Zophobas atratus* | 13 | 2.86 |
|  |  | 14 | 11.43 |
|  |  | 15 | 17.14 |
|  |  | 16 | 25.71 |
|  |  | 17 | 25.71 |
|  |  | 18 | 17.14 |
|  | *Lasiocampa pini* | 4 | 7.14 |
|  |  | 5 | 69.05 |
|  |  | 6 | 19.05 |
|  |  | 7 | 4.76 |

**Table** **S7**: Mean RGR estimates for each species with standard errors ascending ordered within Hemimetabola and Holometabola, respectively.

|  | Order | Species | Instars | | RGR | RGR SE | Weighted |
| --- | --- | --- | --- | --- | --- | --- | --- |
|  |  |  | No | Used |  |  |  |
| Hemimetabola | Phasmatodea | *Dixippus morosus* | 6 | 6 | 0.003 | 0.002 | No |
|  | Blattodea | *Eucorydia yasumatsui* | 10 | 10 | 0.004 | 0.001 | Yes |
|  | Odonata | *Aeshna tuberculifera* | 30 | 29 | 0.011 | 0.002 | No |
|  | Zoraptera | *Zorotypus caudelli* | 5 | 5 | 0.011 | 0.004 | No |
|  | Orthoptera | *Trilophidia conturbata* | 5 | 5 | 0.011 | 0.005 | No |
|  | Odonata | *Leucorrhinia dubia* | 13 | 13 | 0.013 | 0.003 | No |
|  | Odonata | *Nasiaeschna pentacantha* | 15 | 13 | 0.013 | 0.002 | No |
|  | Orthoptera | *Acrida sp.* | 7 | 7 | 0.013 | 0.002 | No |
|  | Orthoptera | *Acrotylus angulatus* | 5 | 5 | 0.014 | 0.002 | No |
|  | Orthoptera | *Acheta domesticus* | 10 | 10 | 0.018 | 0.006 | No |
|  | Orthoptera | *Oedaleus nigrofasciatus* | 5 | 5 | 0.018 | 0.003 | No |
|  | Orthoptera | *Zonocerus elegans* | 5 | 4 | 0.019 | 0.003 | No |
|  | Orthoptera | *Teleogryllus commodus* | 10 | 10 | 0.020 | 0.006 | No |
|  | Orthoptera | *Gryllus assimilis* | 10 | 10 | 0.020 | 0.006 | No |
|  | Orthoptera | *Acanthacris ruficornis* | 7 | 6 | 0.020 | 0.009 | No |
|  | Psocodea | *Pediculus humanus corpor* | 3 | 2 | 0.023 | 0.003 | No |
|  | Orthoptera | *Aiolopus thalassinus* | 5 | 5 | 0.024 | 0.004 | No |
|  | Odonata | *Anax junius* | 13 | 12 | 0.029 | 0.011 | No |
|  | Hemiptera | *Psammotettix alienus* | 5 | 4 | 0.037 | 0.004 | No |
|  | Odonata | *Erythemis simplicicollis* | 13 | 12 | 0.048 | 0.007 | No |
|  | Hemiptera | *Limnogeton fieberi* | 5 | 4 | 0.047 | 0.013 | No |
| Holometabola | Lepidoptera | *Epilobophora sabinata teriolensis* | 5 | 3 | 0.008 | 0.003 | No |
|  | Coleoptera | *Cucujus cinnaberinus* | 6 | 5 | 0.009 | 0.002 | No |
|  | Coleoptera | *Tenebrio molitor* | 20 | 19 | 0.015 | 0.003 | Yes |
|  | Lepidoptera | *Lasiocampa pini* | 7 | 6 | 0.019 | 0.007 | Yes |
|  | Coleoptera | *Zophobas atratus* | 18 | 18 | 0.021 | 0.002 | Yes |
|  | Lepidoptera | *Saturnia pyri* | 6 | 5 | 0.088 | 0.022 | No |
|  | Lepidoptera | *Ephestia kuehniella* | 6 | 6 | 0.096 | 0.012 | No |
|  | Coleoptera | *Melasoma populi* | 4 | 4 | 0.089 | 0.046 | No |
|  | Lepidoptera | *Hyles livornica* | 5 | 3 | 0.111 | 0.063 | No |
|  | Coleoptera | *Tribolium confusum* | 6 | 6 | 0.138 | 0.035 | No |
|  | Coleoptera | *Andrector ruficornis* | 3 | 3 | 0.263 | 0.081 | No |
|  | Diptera | *Lucilia sericata* | 3 | 2 | 1.012 | 0.192 | No |

**Table** **S8**: *Listed is the per species information about the rearing temperature, where the study was conducted (field or laboratory), and whether living specimens or exuviae were measured.*

|  | Order | Species | Laboratory / Field | Temp | living insect |
| --- | --- | --- | --- | --- | --- |
| Hemimetabola | Blattodea | *Eucorydia yasumatsui* | Laboratory | 18-24 | living |
|  | Hemiptera | *Limnogeton fieberi* | Laboratory | 28 | living |
|  |  | *Psammotettix alienus* | Laboratory | >20 | living |
|  | Odonata | *Aeshna tuberculifera* | Laboratory | RT | exuvie and living |
|  |  | *Anax junius* | Laboratory | 10 - 31.6 | exuvie |
|  |  | *Nasiaeschna pentacantha* | Laboratory | RT | exuvie |
|  |  | *Erythemis simplicicollis* | Laboratory | RT | exuvie |
|  |  | *Leucorrhinia dubia* | Field | RT | living |
|  | Orthoptera | *Acanthacris ruficornis* | Laboratory | Africa | living |
|  |  | *Acrida sp.* | Laboratory | Africa | living |
|  |  | *Acrotylus angulatus* | Laboratory | Africa | living |
|  |  | *Aiolopus thalassinus* | Laboratory | Africa | living |
|  |  | *Oedaleus nigrofasciatus* | Laboratory | Africa | living |
|  |  | *Trilophidia conturbata* | Laboratory | Africa | living |
|  |  | *Acheta domesticus* | Laboratory | 25 | living |
|  |  | *Gryllus assimilis* | Laboratory | 25 | living |
|  |  | *Teleogryllus commodus* | Laboratory | 25 | living |
|  |  | *Zonocerus elegans* | Laboratory | Africa | living |
|  | Phasmatodea | *Dixippus morosus* | Laboratory | 20 | living |
|  | Psocodea | *Pediculus humanus corpor* | Laboratory | RT | living |
|  |  | *Pediculus humanus corpor* | Laboratory | RT | living |
|  | Zoraptera | *Zorotypus caudelli* | Laboratory | 26 | living |
| Holometabola | Coleoptera | *Andrector ruficornis* | Laboratory | 28 | living |
|  |  | *Melasoma populi* | Laboratory | RT | living |
|  |  | *Cucujus cinnaberinus* | Laboratory | 22 | living |
|  |  | *Tenebrio molitor* | Laboratory | 25 | living |
|  |  | *Tribolium confusum* | Laboratory | 30 | living |
|  |  | *Zophobas atratus* | Laboratory | 25 | living |
|  | Diptera | *Lucilia sericata* | Laboratory | 25 | living |
|  | Lepidoptera | *Epilobophora sabinata teriolensis* | Field | shady and cool | living |
|  |  | *Lasiocampa pini* | Lab. / Field | RT | living |
|  |  | *Ephestia kuehniella* | Laboratory | 30 | living |
|  |  | *Saturnia pyri* | Laboratory | RT | living |
|  |  | *Hyles livornica* | Laboratory | <20 | living |

**Table S9:** References for the growth data.

|  | | Species | Author | Title | Journal | Vol, No., pp. | Year | Language |
| --- | --- | --- | --- | --- | --- | --- | --- | --- |
| Hemimetabola | Blattodea | *Eucorydia yasumatsui* | Fujita Mari, Ryuichiro Machida | Reproductive biology and postembryonic development of a polyphagid cockroach Eucorydia yasumatsui Asahina (Blattodea: Polyphagidae) | Arthropod Systematics & Phylogeny | 72, 2, 193-211 | 2014 | English |
|  | Hemiptera | *Limnogeton fieberi* | J. Voelker | Untersuchungen zu Ernährung, Fortpflanzungsbiologie und Entwicklung von Limnogeton fieberi Mayr (Belostomatidae, Hemiptera) als Beitrag zur Kenntnis von natürlichen Feinden tropischer Wasserschnecken | Entomologische Mitteilungen aus dem Zoologischesn Staatsinstitut u. Zoologischen Museum Hamburg | 3,60 | 1968 | German |
|  |  | *Psammotettix alienus* | Binari Manurung, Werner Witsack, Egon Fuchs, Silke Mehner | Zur Embryonal-und Larvalentwicklung der Zikade Psammotettix alienus (DAHLBOM, 1851):(Hemiptera, Auchenorrhyncha) | Cicadina | 4, 49-58 | 2001 | German |
|  | Odonata | *Aeshna tuberculifera* | Elsie Lincoln | Growth in Aeshna Tuberculifera (Odonata) | Proceedings of the American Philosophical Society | 83, 5, 589-605 | 1940 | English |
|  |  | *Anax junius* | Philip P. Calvert | The Rates of Growth, Larval Development and Seasonal Distribution of Dragonflies of the Genus Anax (Odonata: Æshnidæ) | Proceedings of the American Philosophical Society | 73, 1, 1-70 | 1934 | English |
|  |  | *Nasiaeschna pentacantha* | S.W. Dunkle | Larval growth in Nasiaeschna pentacantha (Rambur)(Anisoptera: Aeshnidae) | Odonatologica | 14, 1, 29-35 | 1985 | English |
|  |  | *Erythemis simplicicollis* | George H. Bick | Life-history of the dragofly, Erythemis simplicicollis (Say) | Annals of the Entomological Society of America | 34, 1, 215-230 | 1941 | English |
|  |  | *Leucorrhinia dubia* | Fritz Prenn | Aus der Nordtiroler Libellenfauna | Buch; Verlag: K.k.zoologisch-botanische Gesellschaft, Wien |  | 1929 | German |
|  | Orthoptera | *Acanthacris ruficornis* | Julia Chesler | Observations on the biology of some South African Acrididae (Orthoptera) | Transactions of the Royal Entomological Society of London | 87, 14, 313-351 | 1938 | English |
|  |  | *Acrida sp.* |  |  |  |  |  |  |
|  |  | *Acrotylus angulatus* |  |  |  |  |  |  |
|  |  | *Aiolopus thalassinus* |  |  |  |  |  |  |
|  |  | *Oedaleus nigrofasciatus* |  |  |  |  |  |  |
|  |  | *Trilophidia conturbata* |  |  |  |  |  |  |
|  |  | *Zonocerus elegans* |  |  |  |  |  |  |
|  |  | *Acheta domesticus* | Robert Sturm | Längen- und Gewichtsentwicklung der Larven verschiedener Grillenarten (Orthoptera: Gryllidae) vom Zeitpunkt des Ausschlüpfens bis zur Adulthäutung | Linzer biologische Beiträge | 35, 1, 487-498 | 2003 | German |
|  |  | *Gryllus assimilis* |  |  |  |  |  |  |
|  |  | *Teleogryllus commodus* |  |  |  |  |  |  |
|  | Phasmatodea | *Dixippus morosus* | Eidmann | Untersuchungen über Wachstum und Häutung der Insekten | Zeitschrift für Morphologie und Ökologie der Tiere | 2, 3-4, 567-610 | 1924 | German |
|  | Psocodea | *Pediculus humanus corpor* | P.A. Buxton | Studies on the growth of Pediculus (Anoplura) | Parasitology | 30,1, 65-84 | 1938 | English |
|  |  | *Pediculus humanus corpor* | P.A. Buxton | The biology of the body louse (Pediculus humanus corporis: Anoplura) under experimental conditions | Parasitology | 32, 3, 303-312 | 1940 | English |
|  | Zoraptera | *Zorotypus caudelli* | Yuta Mashimo, Rolf G. Beutel, Romano Dallai, Chow-Yang Lee, Ryuichiro Machida | Postembryonic development of the ground louse Zorotypus caudelli Karny (Insecta: Zoraptera: Zorotypidae) | Arthropod Systematics & Phylogeny | 72, 1, 55-71 | 2014 | English |
| Holometabola | Coleoptera | *Andrector ruficornis* | Wolfgang Heyer, Maria Luisa Chiang Lok, Bienvenido Cruz | Zum Einfluss der Temperatur und Wirtspflanze auf die Entwicklung von Andrector ruficornis (Oliv.) (Coleoptera: Chrysomelidae) | Beiträge zur Entomologie | 38, 1, 183-188 | 1988 | German |
|  |  | *Melasoma populi* | A. Willer | Beobachtungen zur Biologie von Melasoma populi L. | Zeitschrift für wissenschaftliche Insektenbiologie | 15, 65-73 | 1919 | German |
|  |  | *Cucujus cinnaberinus* | Ulrich Straka | Zur Biologie des Scharlachkäfers Cucujus cinnaberinus (Scopoli, 1763) | Beiträge zur Entomofaunistik | 8, 11-26 | 2008 | German |
|  |  | *Tenebrio molitor* | Jong Bin Park, Won Ho Choi, Seong Hyun Kim, Hyo Jung Jin, Yeon Soo Han, Yong Seok Lee, Nam Jung Kim | Developmental characteristics of Tenebrio molitor larvae (Coleoptera: Tenebrionidae) in different instars | International Journal of Industrial Entomology | 28, 1, 5-9 | 2014 | English |
|  |  | *Tribolium confusum* | Tom A. Brindley | The Growth and Development of Ephestia Kuehniella Zeller (Lepidoptera) and Tri-Bolium Confusum Duval (Coleoptera) under Controlled Conditions of Temperature and Relative Humidity | Annals of the Entomological Society of America | 23, 4, 741-757 | 1930 | English |
|  |  | *Zophobas atratus* | Sun Young Kim, Hong Geun Kim, Sung Ho Song, Nam Jung Kim | Developmental characteristics of Zophobas atratus (Coleoptera: Tenebrionidae) larvae in different instars | International Journal of Industrial Entomology | 30, 2, 45-49 | 2015 | English |
|  | Diptera | *Lucilia sericata* | Martin Grassberger, Christian Reiter | Effect of temperature on Lucilia sericata (Diptera: Calliphoridae) development with special reference to the isomegalen-and isomorphen-diagram | Forensic Science International | 120, 1-2, 32-36 | 2001 | English |
|  | Lepidoptera | *Epilobophora sabinata teriolensis* | Wilhelm Mack | Die Entwicklung von Nothopteryx (Lobophora) sabinata H.-Schäf.f von teriolensis Kitt | Zeitschrift des Wienerr Entomologen-Vereins | 27, 16-22 | 1942 | German |
|  |  | *Lasiocampa pini* | Karl Eckstein | Beiträge zur Kenntnis des Kiefernspinners Lasiocampa (Gastropacha, Dendrolimus) pini L. | Zoologische Jahrbücher, Abteilung für Systematik, Geographie und Biologie der Tiere | 31, 59-164 | 1911 | German |
|  |  | *Ephestia kuehniella* | Tom A. Brindley | The Growth and Development of Ephestia Kuehniella Zeller (Lepidoptera) and Tri-Bolium Confusum Duval (Coleoptera) under Controlled Conditions of Temperature and Relative Humidity | Annals of the Entomological Society of America | 23, 4, 741-757 | 1930 | English |
|  |  | *Saturnia pyri* | Oliver Eitschberger | Die Biologie und Metamorphose des Wiener Nachtpfauenauges Saturnia pyri ([Denis & Schiffermüller], 1775) | Neue Entomologische Nachrichten | 148-171 | 2010 | German |
|  |  | *Hyles livornica* | M. Gillmer | Ein Beitrag zur Entwicklungsgeschichte von Phryxus livornica, Esp. | Entomologische Zeitschrift | 70-72 | 1904 | German |

**Model description:**

We developed a minimal model of an insect life cycle with a phase of growth of duration *t* during which allocation towards differentiation can occur, followed by a (facultative) phase of pure differentiation, of duration $\Delta$ (Figure 2). Both growth and differentiation saturate, reflecting the existence of an asymptotic size, L_T,_ and a final state of differentiation beyond which further differentiation is impossible, set to 1; both are modeled using the von Bertalanffy growth curve, curves appropriate to describe the two distinct developmental phases up to adulthood (metamorphosis), with reproduction occurring thereafter (appropriate since our model largely proceeds allocation towards reproductive structures, see below). Expected fertility, *F*, is defined as final size modulated by the degree of differentiation that has occurred at the end of life, such that if an organism reaches the final size *L_T_*, and has also achieved maximal differentiation, 1, *F*= *L_T_*^1+1^ *=L_T_*^2^; similar qualitative results are obtained if fitness is defined as the product of growth and differentiation.

To capture the trade-off between growth and differentiation, we assume that the insect has a finite pool of resources that can be allocated either towards the growth rate, termed *r_1_*, the rate of differentiation during growth, termed *r_2_*, or differentiation after growth, termed *r_3_*, with *r_1_+r_2_+r_3_*=1. We explore optimal life histories in terms of allocation towards *r_1_* and *r_2_* and optimal duration of the growth phase, *t*. We define fitness by the net reproductive number, $R_{0}$, obtained by combining fertility, differentiation and survival over the two life phases. Since there is little information to inform the duration of, and difference between allocation to differentiation and survival over the final differentiation phase of life, we simplify this part of the model, assuming that resources allocated towards this part of the life cycle, *r_3_*, capture the proportion of remaining differentiation achieved. The final expression for fitness is:

$$R_{0}=e^{-\mu_{G}t}\left[ L_{T}\left( 1-e^{-\left( k+r_{1} \right)\left( t-v \right)} \right) \right]^{1+\left( 1-e^{-r_{2}\left( t-v \right)} \right)+r_{3}e^{-r_{2}\left( t-v \right)}}$$

The first term $e^{-\mu_{G}t}$, captures survival over the growth phase, the second term is growth in size ($L_{T}\left( 1-e^{-\left( k+r_{1} \right)\left( t-v \right)} \right)$, with asymptotic size $L_{T}$, growth rate $k+r_{1}$, where *k* is the background growth efficiency realized in the absence of resource allocation, and $v$ capturing the hypothetical age at which size would be zero, required to prevent negative growth. Differentiation during the growth phase occurs at rate $r_{2}$, and the proportion of the remainder of differentiation that is possible ($e^{-r_{2}\left( t-v \right)}$) that occurs after the growth phase is set by $r_{3}$. Holometabolous life cycles are reflected by no differentiation occurring during the growth phase, i.e., $r_{2}=0$, and allocation towards differentiation after the growth phase, i.e., $r_{3}>0$.

Choice of von Bertalanffy growth curves. Insect growth curves have been described by various models including the one of von Bertalanffy, a specific case of a generalized logarithmic function (Maino & Kearney 2015). This growth function has been criticized to model optimal size and age at maturity in insects (Day & Taylor 1997). The main critique formulates the need to separate a prematurity period—where no surplus energy is devoted to reproduction—and a post-maturity period, where all energy is devoted to reproduction, a separation a single von Bertalanffy model does not achieve (Day & Taylor 1997). Our model separates such distinct phases of growth and differentiation, as demanded by the criticizing theory of the Bertalanffy, but does it a less absolute way, i.e. growth and differentiation are not completely separated during the first period, also reproduction is an integrated measure of growth and differentiation in our model, combined, in doing so, we are able to use distinct von Bertalanffy models as a base for both periods of growth and development and still not violating the critique for using a single Bertalanffy growth function to model optimal age and size at maturity.

**Table S10: Model parameters and descriptions**

| Symbol | Description |
| --- | --- |
| $L_{T}$ | Asymptotic size |
| $k$ | Rate of growth in size |
| $t$ | Age at the end of the period of growth in size |
| $v$ | Hypothetical age at which size would be zero |
| $\Delta$ | Age at the end of the period of differentiation |
| $\mu_{G}$ | Rate of mortality during the growth phase |
| $F$ | Fertility, defined as the product of growth in size and differentiation at the end of each respective phase ($F=L_{t}D_{\Delta}$) |
| $R_{0}$ | Net reproductive number across both the growth and differentiation phase |
| $r_{1}$ | Allocation of resources towards growth in size |
| $r_{2}$ | Allocation of resources towards differentiation during the growth phase |
| $r_{3}$ | Allocation of resources towards differentiation in the final phase |

References

1.	N. A. Moran, Adaptation and Constraint in the Complex Life Cycles of Animals. *Annual Review of Ecology and Systematics* **2**, 573–600 (1994).

2.	E. E. Werner, Amphibian Metamorphosis: Growth Rate, Predation Risk, and the Optimal Size at Transformation. *The American Naturalist* **128**, 319–341 (1986).

3.	H. ten Brink, A. M. de Roos, U. Dieckmann, The Evolutionary Ecology of Metamorphosis. *The American Naturalist* **193**, E116–E131 (2019).

4.	C. Mora, D. P. Tittensor, S. Adl, A. G. B. Simpson, B. Worm, How Many Species Are There on Earth and in the Ocean? *PLoS Biol* **9**, e1001127 (2011).

5.	B. Misof, S. Liu, K. Meusemann, R. S. Peters, A. Donath, C. Mayer, P. B. Frandsen, J. Ware, T. Flouri, R. G. Beutel, O. Niehuis, M. Petersen, F. Izquierdo-Carrasco, T. Wappler, J. Rust, A. J. Aberer, U. Aspöck, H. Aspöck, D. Bartel, A. Blanke, S. Berger, A. Böhm, T. R. Buckley, B. Calcott, J. Chen, F. Friedrich, M. Fukui, M. Fujita, C. Greve, P. Grobe, S. Gu, Y. Huang, L. S. Jermiin, A. Y. Kawahara, L. Krogmann, M. Kubiak, R. Lanfear, H. Letsch, Y. Li, Z. Li, J. Li, H. Lu, R. Machida, Y. Mashimo, P. Kapli, D. D. McKenna, G. Meng, Y. Nakagaki, J. L. Navarrete-Heredia, M. Ott, Y. Ou, G. Pass, L. Podsiadlowski, H. Pohl, B. M. Von Reumont, K. Schütte, K. Sekiya, S. Shimizu, A. Slipinski, A. Stamatakis, W. Song, X. Su, N. U. Szucsich, M. Tan, X. Tan, M. Tang, J. Tang, G. Timelthaler, S. Tomizuka, M. Trautwein, X. Tong, T. Uchifune, M. G. Walzl, B. M. Wiegmann, J. Wilbrandt, B. Wipfler, T. K. F. Wong, Q. Wu, G. Wu, Y. Xie, S. Yang, Q. Yang, D. K. Yeates, K. Yoshizawa, Q. Zhang, R. Zhang, W. Zhang, Y. Zhang, J. Zhao, C. Zhou, L. Zhou, T. Ziesmann, S. Zou, Y. Li, X. Xu, Y. Zhang, H. Yang, J. Wang, J. Wang, K. M. Kjer, X. Zhou, Phylogenomics resolves the timing and pattern of insect evolution. *Science* **346**, 763–767 (2014).

6.	B. Wang, C. Xu, E. A. Jarzembowski, Ecological radiations of insects in the Mesozoic. *Trends in Ecology & Evolution*, S0169534722000441 (2022).

7.	D. A. Grimaldi, M. Engel, *The Evolution of the Insects* (Cambridge University Press, Cambridge, 2005).

8.	J. Rolff, P. R. Johnston, S. Reynolds, Complete metamorphosis of insects. *Phil. Trans. R. Soc. B* **374**, 20190063 (2019).

9.	M. J. R. Hall, D. Martín-Vega, Visualization of insect metamorphosis. *Phil. Trans. R. Soc. B* **374**, 20190071 (2019).

10.	X. Bellés, *Insect Metamorphosis: From Natural History to Regulation of Development and Evolution* (Academic Press, London, 2020).

11.	S. Aldaz, L. Escudero, Imaginal discs. *Current Biology*, R429–R431.

12.	R. Chapman, *The Insects - Structure and Function* (Cambridge University Press, Cambridge, 2013).

13.	F. Sehnal, Morphology of Insect Development.

14.	H. E. Hinton, THE ORIGIN AND FUNCTION OF THE PUPAL STAGE. *Proceedings of the Royal Entomological Society of London. Series A, General Entomology* **38**, 77–85 (1963).

15.	J. W. Truman, L. M. Riddiford, The evolution of insect metamorphosis: a developmental and endocrine view. *Phil. Trans. R. Soc. B* **374**, 20190070 (2019).

16.	S. Reynolds, Cooking up the perfect insect: Aristotle’s transformational idea about the complete metamorphosis of insects. *Phil. Trans. R. Soc. B* **374**, 20190074 (2019).

17.	M. A. Albecker, L. G. E. Wilkins, S. A. Krueger-Hadfield, S. M. Bashevkin, M. W. Hahn, M. P. Hare, H. K. Kindsvater, M. A. Sewell, K. E. Lotterhos, A. M. Reitzel, Does a complex life cycle affect adaptation to environmental change? Genome-informed insights for characterizing selection across complex life cycle. *Proc. R. Soc. B.* **288**, 20212122 (2021).

18.	J. D. Arendt, Adaptive Intrinsic Growth Rates: An Integration Across Taxa. *The Quarterly Review of Biology* **72**, 149–177 (1997).

19.	D. A. Herms, W. J. Mattson, The Dilemma of Plants: To Grow or Defend. *The Quarterly Review of Biology* **67**, 283–335 (1992).

20.	J. Rolff, F. V. de Meutter, R. Stoks, Time Constraints Decouple Age and Size at Maturity and Physiological Traits. 7.

21.	C. M. Dmitriew, The evolution of growth trajectories: what limits growth rate? *Biological Reviews* **86**, 97–116 (2011).

22.	L. Rowe, D. Ludwig, Size and timing of metamorphosis in complex life cycles: times constraints and variation. *Ecology*, 413–427.

23.	L. D. Mueller, Density-Dependent Population Growth and Natural Selection in Food-Limited Environments: The Drosophila Model. *The American Naturalist* **132**, 786–809 (1988).

24.	K. Spitze, *CHAOBORUS* PREDATION AND LIFE‐HISTORY EVOLUTION IN *DAPHNIA PULEX* : TEMPORAL PATTERN OF POPULATION DIVERSITY, FITNESS, AND MEAN LIFE HISTORY. *Evolution* **45**, 82–92 (1991).

25.	P. T. Rühr, T. van de Kamp, T. Faragó, J. U. Hammel, F. Wilde, E. Borisova, C. Edel, M. Frenzel, T. Baumbach, A. Blanke, Juvenile ecology drives adult morphology in two insect orders. *Proc. R. Soc. B.* **288**, 20210616 (2021).

26.	C. J. E. Metcalf, B. Koskella, Protective microbiomes can limit the evolution of host pathogen defense. *Evolution Letters* **3**, 534–543 (2019).

27.	I. J. Banks, W. T. Gibson, M. M. Cameron, Growth rates of black soldier fly larvae fed on fresh human faeces and their implication for improving sanitation. *Tropical Med Int Health* **19**, 14–22 (2014).

28.	P. T. Smiseth, C. T. Darwell, A. J. Moore, Partial begging: an empirical model for the early evolution of offspring signalling. *Proc. R. Soc. Lond. B* **270**, 1773–1777 (2003).

29.	C. Jaspers, R. R. Hopcroft, T. Kiørboe, F. Lombard, Á. López-Urrutia, J. D. Everett, A. J. Richardson, Gelatinous larvacean zooplankton can enhance trophic transfer and carbon sequestration. *Trends in Ecology & Evolution* **38**, 980–993 (2023).

30.	M. Denoël, P. Joly, H. H. Whiteman, Evolutionary ecology of facultative paedomorphosis in newts and salamanders. *Biological Reviews* **80**, 663–671 (2005).

31.	D. P. Mcmahon, A. Hayward, Why grow up? A perspective on insect strategies to avoid metamorphosis. *Ecological Entomology* **41**, 505–515 (2016).

32.	J. W. Truman, L. M. Riddiford, Endocrine Insights into the Evolution of Metamorphosis in Insects. *Annu. Rev. Entomol.* **47**, 467–500 (2002).

33.	A. Berlese, Intorno alle metamorfosi degli insetti. *Redia* **9**, 121–136.

34.	S. E. Reynolds, A transcription factor that enables metamorphosis. *Proc. Natl. Acad. Sci. U.S.A.* **119**, e2204972119 (2022).

35.	J. W. Truman, L. M. Riddiford, *Chinmo* is the larval member of the molecular trinity that directs *Drosophila* metamorphosis. *Proc. Natl. Acad. Sci. U.S.A.* **119**, e2201071119 (2022).

36.	A. Fernandez-Nicolas, G. Machaj, A. Ventos-Alfonso, V. Pagone, T. Minemura, T. Ohde, T. Daimon, G. Ylla, X. Belles, Reduction of embryonic *E93* expression as a hypothetical driver of the evolution of insect metamorphosis. *Proc. Natl. Acad. Sci. U.S.A.* **120**, e2216640120 (2023).

37.	J. W. Truman, The Evolution of Insect Metamorphosis. *Current Biology* **29**, R1252–R1268 (2019).

38.	M. Jindra, Where did the pupa come from? The timing of juvenile hormone signalling supports homology between stages of hemimetabolous and holometabolous insects. *Phil. Trans. R. Soc. B* **374**, 20190064 (2019).

39.	S. Stankowski, Z. B. Zagrodzka, M. D. Garlovsky, A. Pal, D. Shipilina, D. G. Castillo, H. Lifchitz, A. L. Moan, E. Leder, J. Reeve, K. Johannesson, A. M. Westram, R. K. Butlin, The genetic basis of a recent transition to live-bearing in marine snails. (2024).

40.	C. Manthey, P. R. Johnston, S. Nakagawa, J. Rolff, Complete metamorphosis and microbiota turnover in insects. *Molecular Ecology*, mec.16673 (2022).

41.	T. J. Hammer, N. A. Moran, Links between metamorphosis and symbiosis in holometabolous insects. *Phil. Trans. R. Soc. B* **374**, 20190068 (2019).

42.	H. Hinton, On the origin and function of the pupal stage. *Transactions of the Royal Entomological Society* **99**, 395–409.

43.	W. D. Hamilton, “Funeral Feasts: Evolution and diversity under barke” in *Narrow Roads of Gene Land* (WH Freeman, New York, NY, ed. 1, 1996)vol. 1, pp. 387–422.

44.	R. S. Peters, K. Meusemann, M. Petersen, C. Mayer, J. Wilbrandt, T. Ziesmann, A. Donath, K. M. Kjer, U. Aspöck, H. Aspöck, A. Aberer, A. Stamatakis, F. Friedrich, F. Hünefeld, O. Niehuis, R. G. Beutel, B. Misof, The evolutionary history of holometabolous insects inferred from transcriptome-based phylogeny and comprehensive morphological data. *BMC Evol Biol* **14**, 52 (2014).

45.	V. Wigglesworth, *The Physiology of Insect Metamorphosis* (Cambridge University Press, Cambridge, 1954)vol. 1 of *Cambridge Monographs in experimental Biology I*.

46.	S. Fellous, B. P. Lazzaro, Potential for evolutionary coupling and decoupling of larval and adult immune gene expression. *Molecular Ecology* **20**, 1558–1567 (2011).

47.	V. Loeschcke, R. A. Krebs, SELECTION FOR HEAT-SHOCK RESISTANCE IN LARVAL AND IN ADULT *DROSOPHILA BUZZATII* : COMPARING DIRECT AND INDIRECT RESPONSES. *Evolution* **50**, 2354–2359 (1996).

48.	D. B. Nicholson, A. J. Ross, P. J. Mayhew, Fossil evidence for key innovations in the evolution of insect diversity. *Proc. R. Soc. B.* **281**, 20141823 (2014).

49.	A. S. Yang, Modularity, evolvability, and adaptive radiations: a comparison of the hemi‐ and holometabolous insects. *Evolution and Development* **3**, 59–72 (2001).

50.	T. Tammaru, S. Nylin, K. Ruohomäki, K. Gotthard, Compensatory responses in lepidopteran larvae: a test of growth rate maximisation. *Oikos* **107**, 352–362 (2004).

51.	R Core Team, R: A language and environment for statistical	computing. R Foundation for Statistical Computing, Vienna, Austria. ., (2021); URL https://www.R-project.org/.

52.	P. Dutilleul, P. Legendre, Spatial Heterogeneity against Heteroscedasticity: An Ecological Paradigm versus a Statistical Concept. *Oikos* **66**, 152 (1993).

53.	W. Fox, J S., *An R Companion to Applied Regression, Third Edition. Sage, Thousand Oaks CA* (Sage, Thousand Oaks, CA, 2019; https://socialsciences.mcmaster.ca/jfox/Books/Companion/.).

54.	A. GRAFENt, The Phylogenetic Regression. (2023).

55.	O. Cinar, S. Nakagawa, W. Viechtbauer, Phylogenetic multilevel meta‐analysis: A simulation study on the importance of modelling the phylogeny. *Methods Ecol Evol* **13**, 383–395 (2022).

56.	S. Nakagawa, M. Lagisz, R. E. O’Dea, J. Rutkowska, Y. Yang, D. W. A. Noble, A. M. Senior, The orchard plot: Cultivating a forest plot for use in ecology, evolution, and beyond.

57.	S. Nakagawa, H. Schielzeth, A general and simple method for obtaining *R* ^2^ from generalized linear mixed‐effects models. *Methods Ecol Evol* **4**, 133–142 (2013).

R-Scripts

Phylogenetic model

library(tidyverse)

library(ape)

library(metafor)

library(orchaRd)

library(ape)

library(phytools)

library(tidyverse)

#############################################

\# Import DF

#############################################

ALL_GR_summarySE <- read.csv("TableS7.csv", header = TRUE, dec=",", sep="\t", na.strings = "na")

#############################################

\# Make Phylogeny

#############################################

text.string<- "((Odonata(Aeshnidae(Aeshna_tuberculifera,Anax_junius,Nasiaeschna_pentacantha), Libellulidae(Erythemis_simplicicollis,Leucorrhinia_dubia))),(((Zoraptera(Zorotypidae(Zorotypus_caudelli))),((Orthoptera((Acrididae(Cyrtacanthacridinae(Acanthacris_ruficornis), (Acridinae(Acrida_sp), Oedipodinae(Trilophidia_conturbata, Oedaleus_nigrofasciatus, Aiolopus_thalassinus, Acrotylus_angulatus))), Pyrgomorphidae(Zonocerus_elegans)), Gryllidae(Acheta_domesticus, Gryllus_assimilis, Teleogryllus_commodus))),((Phasmatodea(Phasmatidae(Dixippus_morosus))),(Blattodea(Polyphagidae(Eucorydia_yasumatsui)))))),((Hemiptera(Belostomatidae(Limnogeton_fieberi),Cicadellidae_Jassidae(Psammotettix_alienus))),((Psocodea(Pediculidae(Pediculus_humanus_corpor))),((Coleoptera(Tenebrionidae(Tenebrio_molitor, Tribolium_confusum, Zophobas_atratus), (Chrysomelinae(Melasoma_populi, Andrector_ruficornis), Cucujidae(Cucujus_cinnaberinus)))),((Lepdioptera(Pyralidae(Ephestia_kuehniella), (Geometridae(Epilobophora_sabinata_teriolensis), (Lasiocampidae(Lasiocampa_pini), (Saturniidae(Saturnia_pyri), Sphingidae(Hyles_livornica)))))),(Diptera(Caliphoridae(Lucilia_sericata)))))))));"

vert.tree<-ape::read.tree(text=text.string)

plotTree(vert.tree,offset=2)

#############################################

\# Generate Grafen's Branch Lengths

#############################################

\# turning into phylogenetic correlation matrix using Grafen's method

tree <- compute.brlen(vert.tree) # Generate branch lengths; method = "Grafen" --> default

cor_tree <- vcv(tree,corr = T) # Generate phylogenetic variance-covariance matrix

######################################################

\# Add observational level data

######################################################

ALL_GR_summarySE$obs <- factor(1:dim(ALL_GR_summarySE)[1])

#############################################

\# Heteroscedasticity test

#############################################

library(car)

lmMod <- lm(ALL_GR_summarySE$Perc ~ ALL_GR_summarySE$Hemi_Holo)

par(mfrow=c(2,2))

plot(lmMod)

car::ncvTest(lmMod)

#############################################

\# Linear Regression

#############################################

\# Phylogenetic meta-analysis via Multivariate/Multilevel Linear (Mixed-Effects) Models

\# Effect of phylogeny:

model0 <- rma.mv(yi = RGR, V = RGR_se^2, mod = ~ 1,

random = list(~1|Species,

~1|obs),

R = list(Species_Tree = cor_tree),

data = ALL_GR_summarySE,

test = "t"

)

i2_ml(model0)

\# fit the meta-regression model without the intercept for the orchard plot and for the means of 2 groups [summary(model)]

model1 <- rma.mv(yi = Perc, V = Perc_se^2, mod = ~ Hemi_Holo - 1,

random = list(~1|Species,

~1|obs),

R = list(Species_Tree = cor_tree),

data = ALL_GR_summarySE,

test = "t"

)

\# looking at contrast between 2 groups

model1b <- rma.mv(yi = RGR, V = RGR_se^2, mod = ~ 1+ Hemi_Holo,

random = list(~1|Species,

~1|obs),

R = list(Species_Tree = cor_tree),

data = ALL_GR_summarySE,

test = "t"

)

r2_ml(model1b)

orchard_plot(model1, mod = "Hemi_Holo", xlab = "Percentage Growth")

\# modeling heteroscadasticty

model2b <- rma.mv(yi = log(RGR), V = RGR_se^2, mod = ~ Hemi_Holo -1,

random = list(~ Hemi_Holo|Species,

~1|obs),

rho = 0,

struct = "HCS",

R = list(Species_Tree = cor_tree),

data = ALL_GR_summarySE,

test = "t"

)

\# calculate the ‘total’ I^2 along with I^2 for each level

\# describes the percentage of variation across studies/samples that is due to heterogeneity

\# heterogeneity of the distribution of the residuals

i2_ml(model0)

\# calculate marginal and conditional R^2 for mixed models

\# measures the proportion of variation in the response that can be attributed to the predictor

r2_ml(model1b)

##################

\# Orchard_Plot

##################

Orchard_RGR <- orchard_plot_2(model2b, mod = "Hemi_Holo", xlab = "Relative Growth Rate \n [Log scale]", angle=45) + theme( axis.text.y=element_text(size=25, colour="black"), axis.text.x=element_text(size=20, colour="grey20"),axis.title.x = element_text(size = 22, margin = margin(t = 20, r = 0, b = 30, l = 0)), legend.text = element_text(size = 10), legend.title = element_text(size = 12, colour="grey20")) + theme(legend.position="bottom")+ scale_fill_manual(values = c("#D55E00","#0072B2"))+ scale_color_manual(values = c("#D55E00","#0072B2"))+ theme(plot.title = element_text(size=25))#+xlim(0.75,2.5)

png("../Orchard_RGR.png", width = 1.25*480, height = 1*480)

Orchard_RGR

dev.off()

Evolutionary model

#################################################################################################################

\## Code to model a Hemi/Holometabolous life-cycle, with resource allocation between growth and differentiation ##

#################################################################################################################

\## von Bertalannffy growth model

growth <- function(Lt,k,t,v,r1) {

Lt*(1-exp(-(k+r1)*(t-v)))

}

\## von Bertalannffy differentiation model, for differentiation occurring during growth (maps to Figure 2)

change1 <- function(d,t,v,r1,r2,r3) {

(1-exp(-(d+r2)*(t-v)))

}

\## differentiation model, simplified way of capturing differentiation occurring after growth

change2 <- function(d,t,v,r1,r2,r3) {

r3*(1-change1(d=d,t=t,v=v,r1=r1,r2=r2,r3=r3))

}

\## define R0 for a holometab life cycle

R0 <- function(Lt,k,d,t,v,r1,r2,r3, mu1) {

exp(-mu1*(t-v))*growth(Lt=Lt,k=k,t=t,v=v,r1=r1)^(1+change1(d=d,t=t,v=v,r1=r1,r2=r2,r3=r3)+change2(d=d,t=t,v=v,r1=r1,r2=r2,r3=r3))

###exp(-mu1*(t-v))*growth(Lt=Lt,k=k,t=t,v=v,r1=r1)*(1+change1(d=d,t=t,v=v,r1=r1,r2=r2,r3=r3)+change2(d=d,t=t,v=v,r1=r1,r2=r2,r3=r3))

}

\## define R0 for a hemimetab life cycle(i.e., r3 necessarily 0)

R0hemi <- function(Lt,k,d,t,v,r1,r2,r3, mu1) {

exp(-mu1*(t-v))*growth(Lt=Lt,k=k,t=t,v=v,r1=r1)^(1+change1(d=d,t=t,v=v,r1=r1,r2=r2,r3=r3))

###exp(-mu1*(t-v))*growth(Lt=Lt,k=k,t=t,v=v,r1=r1)*(1+change1(d=d,t=t,v=v,r1=r1,r2=r2,r3=r3))

}

\##### 1. Illustrate broad patterns ############################################

\## Choose some parameters

Lt.=10

k.=2

d.=2

t.=10

v.=1

r1.=0.2;r2.=0.2;r3.=0.6;

mu1.=0.2

\## Loop across different durations of the growth phase, calculate and then plot out the fitness landscape

test.t <- seq(1,5,length=100)

R0test.hemi <- R0test <- rep(NA,length(test.t))

for (j in 1:length(test.t)) {

R0test[j] <- R0(Lt=Lt.,k=k.,d=d.,t=test.t[j],v=v.,r1=r1.,r2=r2.,r3=r3., mu1=mu1.)

R0test.hemi[j] <- R0hemi(Lt=Lt.,k=k.,d=d.,t=test.t[j],v=v.,r1=r1.,r2=r2.,r3=r3., mu1=mu1.)

}

plot(test.t,R0test, type="l", xlab="end of growth", ylab=expression(R[0]))

points(test.t,R0test.hemi,type="l",col=2,lty=3)

abline(v=c(test.t[R0test==max(R0test)],test.t[R0test.hemi==max(R0test.hemi)]), col=c(1,2), lty=c(1,3))

\## Start at optimal life history for t from the above, and explore R0 for r1 and r2; with assumption r1 + r2 + r3 = R = 1

t. <- test.t[R0test==max(R0test)]

R. <- 1	#total resource pool

r1test <- seq(0,R.,length=100)

r2test <- seq(0,R.,length=100)

R0.store <- matrix(NA,length(r1test),length(r2test))

for (j in 1:length(r1test)) {

for (k in 1:length(r2test)) {

if ((r1test[j]+r2test[k])>R.) next()

R0.store[j,k] <- R0(Lt=Lt.,k=k.,d=d.,t=t.,v=v.,

r1=r1test[j],r2=min(r2test[k],R.-r1test[j]),r3=R.-r1test[j]-r2test[k], mu1=mu1.)

}}

image(r1test,r2test,exp(R0.store), xlab=expression("Resources to growth, "*r[1]), ylab=expression("Resources to differentiation, "*r[2]))

contour(r1test,r2test,R0.store,add=TRUE)

opt <- which(R0.store==max(R0.store,na.rm=T), arr.ind=T)

r1 <- r1test[opt[1,1]]

r2 <- r2test[opt[1,2]]

r3 <- R.-r1-r2

points(r1,r2,pch=19)

print(c(r1,r2,r3))

\##### 2. Function to identify optima ############################################

\## Function to optimize t, r1, r2 assuming that r1+r2+r3=R

findOpt <- function(par,index=1:3,params=c(5,1,0),Lt=5,k=2,d=1,mu1=0.1,R=1,v=1) {

params[index] <- par

t <- params[1]

r1 <- params[2];

r2 <- params[3];

rc <- (R0(Lt=Lt,k=k,d=d,t=t,v=v,r1=r1,r2=min(r2,R-r1),r3=R-r1-r2, mu1=mu1))

return(-rc)

}

\## Function to optimize t, r1, r2 assuming that r3=0

findOptHemi <- function(par,index=1:2,params=c(5,1,0),Lt=5,k=2,d=1,mu1=0.1,R=1,v=1) {

params[index] <- par

t <- params[1]

r1 <- params[2];

r2 <- R-r1

rc <- (R0hemi(Lt=Lt,k=k,d=d,t=t,v=v,r1=r1,r2=min(r2,R-r1),r3=0, mu1=mu1))

return(-rc)

}

\## Function to optimize t, r1, r2 assuming that r1+r2+r3=R but using a loop rather than the optim function

findOptBigLoop <- function(par,Lt=5,k=2,d=1,mu1=0.1,R=1,v=1,nest=30, do.plot=FALSE) {

t <- par[1]

r1 <- par[2];

r2 <- par[3];

test.t <- seq(max(t-2,1.001),t+2,length=nest)

test.r <- seq(0,1,length=nest)

rc <- array(dim=c(nest,nest,nest))

#print(k)

for (i in 1:nest) {

for (z in 1:nest) {

for (l in 1:nest) {

if ((test.r[z]+test.r[l])>1) next()

rc[i,z,l] <- (R0(Lt=Lt,k=k,d=d,t=test.t[i],v=v,r1=test.r[z],r2=test.r[l],r3=R-test.r[z]-test.r[l], mu1=mu1))

}}}

opt <- which(rc==max(rc,na.rm=TRUE), arr.ind=TRUE)

#create format that looks like what optim returns

tmp <- list()

tmp$par <- c(test.t[opt[1,1]],test.r[opt[1,2]],test.r[opt[1,3]])

tmp$value <- -max(rc,na.rm=TRUE)

if (do.plot) {

image(test.r,test.r,rc[opt[1,1],,], xlab="resources to growth", ylab="resources to differentiate")

contour(test.r,test.r,rc[opt[1,1],,],add=TRUE)

points(tmp$par[2],tmp$par[3],pch=19)

}

return(tmp)

}

\## Sanity check that not missing something by order that optimise the different allocation parameters

findOptBigLoopReverse <- function(par,Lt=5,k=2,d=1,mu1=0.1,R=1,v=1,nest=30, do.plot=FALSE) {

t <- par[1]

r1 <- par[2];

r3 <- par[3];

test.t <- seq(max(t-2,1.001),t+2,length=nest)

test.r <- seq(0,1,length=nest)

rc <- array(dim=c(nest,nest,nest))

#print(k)

for (i in 1:nest) {

for (z in 1:nest) {

for (l in 1:nest) {

if ((test.r[z]+test.r[l])>1) next()

rc[i,z,l] <- (R0(Lt=Lt,k=k,d=d,t=test.t[i],v=v,r1=test.r[z],r2=R-test.r[z]-test.r[l],r3=test.r[l], mu1=mu1))

}}}

opt <- which(rc==max(rc,na.rm=TRUE), arr.ind=TRUE)

#create format that looks like what optim returns

tmp <- list()

tmp$par <- c(test.t[opt[1,1]],test.r[opt[1,2]],R-test.r[opt[1,2]]-test.r[opt[1,3]])

tmp$value <- -max(rc,na.rm=TRUE)

if (do.plot) {

image(test.r,test.r,rc[opt[1,1],,], xlab="resources to growth", ylab="resources to differentiate")

contour(test.r,test.r,rc[opt[1,1],,],add=TRUE)

points(tmp$par[2],tmp$par[3],pch=19)

}

return(tmp)

}

\## sanity check

#tmp <- optim(par=c(5,1,0), findOpt,Lt=5,k=0.001,d=0,mu1=0.1,v=1,R=1, method="L-BFGS-B",lower=c(1.001,0,0),upper=c(30,1,1))

#tmp1 <- findOptBigLoop(par=c(5,1,0), Lt=5,k=0.001,d=0,mu1=0.1,v=1,R=1,nest=30, do.plot=TRUE)

#tmp2 <- findOptBigLoopReverse(par=c(5,1,0), Lt=5,k=0.001,d=0,mu1=0.1,v=1,R=1,nest=30, do.plot=TRUE)

\## check optimize just one parameter...

#R0(Lt=5,k=0.001,d=0,t=tmp1$par[1],v=1,r1=tmp1$par[2],r2=tmp1$par[3],r3=1-tmp1$par[2]-tmp1$par[3],mu1=0.1)

#tmp <- optim(par=4.1,index=1,params=tmp1$par, findOpt,Lt=5,k=0.001,d=0,mu1=0.1,v=1,R=1, method="L-BFGS-B",lower=c(1.001,0,0),upper=c(30,1,1))

\##### 3. Function to identify when to switch to holometabolous ############################################

\## Function to find the optimal switch point

findKswitch <- function(d=0.1,mu1=0.1,R=1,Lt=5, ntest=40,nest=50,do.plot=FALSE, ktest=c(seq(0.01,1.2,length=ntest-1),1.5)) {

#storage; index 1 indicates hemimetab; and is stored first in the R0 matrix...

storePar1 <- storePar <- matrix(NA,ntest,3)

storeR0 <- matrix(NA,ntest,2)

#loop

for (j in 1:ntest) {

##HEMIMETABOLOUS - start and use as starting parameters

tmp1 <- optim(par=c(4,1), index=1:2,params=c(4,1,0),

findOptHemi,Lt=Lt,k=ktest[j],d=d,mu1=mu1,R=R,v=1, method="L-BFGS-B",

lower=c(1.001,0,0),upper=c(30,1,1))

tmp1x <- tmp1

#jiggle to ensure that have optimal

for (xx in 1:10) {

tmp1x <- optim(par=pmin(pmax(rnorm(2,tmp1$par,pmax(tmp1$par/5,0.1)),c(1.001,0)),c(30,1)),

index=1:2,params=tmp1$par,

findOptHemi,Lt=Lt,k=ktest[j],d=d,mu1=mu1,R=R,v=1, method="L-BFGS-B",

lower=c(1.001,0),upper=c(30,1))

if (tmp1x$value<tmp1$value) tmp1 <- tmp1x

}

storePar1[j,] <- c(tmp1$par,0)

storePar1[j,3] <- 1-storePar1[j,2]

##HOLMETABOLOUS

tmp <- findOptBigLoop(par=storePar1[j,1:3],

Lt=Lt,k=ktest[j],d=d,mu1=mu1,R=R,v=1,nest=nest,do.plot=do.plot)

tmpx <- optim(par=tmp$par,findOpt,

index=1:3,params=tmp$par,

Lt=Lt,k=ktest[j],d=d,mu1=mu1,R=R,v=1, method="L-BFGS-B",

lower=c(1.001,0,0),upper=c(30,1,1))

if (tmpx$value<tmp$value) tmp <- tmpx

#jiggle to ensure

for (xx in 1:10) {

tmpx <- optim(par=pmin(pmax(rnorm(3,tmp$par,pmax(tmp$par/5,0.1)),c(1.001,0,0)),c(30,1,1)),

findOpt,Lt=Lt,k=ktest[j],d=d,mu1=mu1,R=R,v=1, method="L-BFGS-B",

lower=c(1.001,0,0),upper=c(30,1,1))

if (tmpx$value<tmp$value) {tmp <- tmpx; print("improve")}

}

\## Put into storage matrix

storePar[j,] <- tmp$par

storePar[j,3] <- min(storePar[j,3],1-storePar[j,2])

\## Store R0

storeR0[j,] <- c(tmp1$value,tmp$value)

}

#find switch point and gradient

if (sum(storePar[,3]<1e-4)>0) {

switch.point <- range(which(storePar[,3]<1e-4,arr.ind=TRUE))

if (min(switch.point)==1) {

switch.index <- max(switch.point)

switch.point <- ktest[max(switch.point)]

kgradient <- storePar[min(switch.index+1,ntest),2]/(ktest[2]-ktest[1])

} else {

switch.index <- (switch.point[1])

switch.point <- ktest[switch.point[1]]

kgradient <- storePar[max(switch.index-1,1),2]/(ktest[2]-ktest[1])

}

} else {

switch.point <- 0; kgradient <- NA

}

return(list(switch.point=switch.point,switch.index=switch.index,

kgradient=kgradient,storePar=storePar,storePar1=storePar1,storeR0=storeR0,ktest=ktest))

}

\## function to plot this out, showing different parameter optima, and the switch point across a gradient of k.

makePlot.k.impact<-function(k.impact){

#[more visually applealing to put the vertical line right before the switch point]

loc.vert.line <- max(k.impact$switch.index-1,1)

vert.line <- k.impact$ktest[max(k.impact$switch.index-1,1)]

par(mfrow=c(1,4))

plot(k.impact$ktest,k.impact$storePar[,1],

xlab=expression("Growth rate, "*italic(k)), ylab=expression("Optimal duration of the growth phase, "*italic(t)),

type="l", ylim=range(c(k.impact$storePar[,1],k.impact$storePar1[,1])),lwd=2)

points(k.impact$ktest,k.impact$storePar1[,1], type="l",lty=3, col="darkgrey",lwd=2)

if (k.impact$switch.point>0) abline(v=vert.line, col=4,lty=1)

points(k.impact$ktest[loc.vert.line],k.impact$storePar[loc.vert.line,1],pch=19)

plot(k.impact$ktest,k.impact$storePar[,2],

xlab=expression("Growth rate, "*italic(k)), ylab=expression("Optimal growth allocation, "*r[1]), type="l", ylim=c(0,1),lwd=2)

points(k.impact$ktest,k.impact$storePar1[,2],type="l", lty=3, col="darkgrey",lwd=2)

if (k.impact$switch.point>0) abline(v=vert.line, col=4,lty=1)

points(k.impact$ktest[loc.vert.line],k.impact$storePar[loc.vert.line,2],pch=19)

plot(k.impact$ktest,k.impact$storePar[,3],

xlab=expression("Growth rate, "*italic(k)), ylab=expression("Optimal growth phase allocation to diff. "*r[2]), type="l", ylim=c(0,1),lwd=2)

points(k.impact$ktest,k.impact$storePar1[,3],type="l", lty=3, col="darkgrey",lwd=2)

if (k.impact$switch.point>0) abline(v=vert.line, col=4,lty=1)

points(k.impact$ktest[loc.vert.line],k.impact$storePar[loc.vert.line,3],pch=19)

plot(k.impact$ktest,pmax(1-k.impact$storePar[,2]-k.impact$storePar[loc.vert.line,3],0),

xlab=expression("Growth rate, "*italic(k)), ylab=expression("Optimal allocation to differentiate post-growth, "*r[3]), type="l", ylim=c(0,1),lwd=2)

if (k.impact$switch.point>0) abline(v=vert.line, col=4,lty=1)

points(k.impact$ktest[loc.vert.line],pmax(1-k.impact$storePar[loc.vert.line,2]-k.impact$storePar[loc.vert.line,3],0),pch=19)

}

\## explore the different optima under low mortality

aa <- findKswitch(d=0,mu1=0.01,R=1,Lt=5, ntest=30,nest=30)

makePlot.k.impact(aa)

\## explore the different optima under high mortality - note r3 should be 1 but is so high makes no difference

aa1 <- findKswitch(d=0,mu1=0.5,R=1,Lt=5, ntest=30,nest=30)

makePlot.k.impact(aa1)

######################################################################################################################################################

#choose a k and then look at fitness across a focal parameter with other parameters set at their optimal, to unpack patterns and compare low and high effects

\## For parameter t ##########################

\### 1. for low mortality

chs.k <- 20

tst.t <- seq(1,9,length=100)

store.R0s <- matrix(NA,length(tst.t),2)

for (jx in 1:length(tst.t)) {

store.R0s[jx,1] <- R0(Lt=5,k=aa$ktest[chs.k],d=0,t=tst.t[jx],v=1,r1=aa$storePar[chs.k,2],r2=aa$storePar[chs.k,3],r3=1-sum(aa$storePar[chs.k,2:3]), mu1=0.01)

store.R0s[jx,2] <- R0hemi(Lt=5,k=aa$ktest[chs.k],d=0,t=tst.t[jx],v=1,r1=aa$storePar1[chs.k,2],r2=aa$storePar1[chs.k,3],r3=0, mu1=0.01)

}

\### 2. for high mortality

store.R0s1 <- matrix(NA,length(tst.t),2)

for (jx in 1:length(tst.t)) {

store.R0s1[jx,1] <- R0(Lt=5,k=aa1$ktest[chs.k],d=0,t=tst.t[jx],v=1,r1=aa1$storePar[chs.k,2],r2=aa1$storePar[chs.k,3],r3=1-sum(aa1$storePar[chs.k,2:3]), mu1=0.5)

store.R0s1[jx,2] <- R0hemi(Lt=5,k=aa1$ktest[chs.k],d=0,t=tst.t[jx],v=1,r1=aa1$storePar1[chs.k,2],r2=aa1$storePar1[chs.k,3],r3=0, mu1=0.5)

}

par(mfrow=c(1,1),bty="l")

plot(tst.t, store.R0s[,1], type="l", xlab="t", ylab=expression(R[0]), ylim=range(store.R0s,c(store.R0s1),na.rm=TRUE),lwd=1.5)

points(tst.t,store.R0s[,2], type="l", col="grey",lty=3,lwd=1.5)

#abline(v=tst.t[store.R0s[,1]==max(store.R0s[,1])])

points(tst.t[store.R0s[,1]==max(store.R0s[,1])],max(store.R0s[,1]),pch=19,cex=1.2,col="black")

#abline(v=tst.t[store.R0s[,2]==max(store.R0s[,2])])

points(tst.t[store.R0s[,2]==max(store.R0s[,2])],max(store.R0s[,2]),pch=19,cex=1.2,col="grey")

#plot(tst.t, store.R0s1[,1], type="l", xlab="t", ylab=expression(R[0]), ylim=range(store.R0s1,na.rm=TRUE),lwd=1.5)

points(tst.t,store.R0s1[,1], type="l", col="black",lty=1,lwd=1.5)

points(tst.t,store.R0s1[,2], type="l", col="grey",lty=3,lwd=1.5)

#abline(v=tst.t[store.R0s1[,1]==max(store.R0s1[,1])])

points(tst.t[store.R0s1[,1]==max(store.R0s1[,1])],max(store.R0s1[,1]),pch=19,cex=1.2,col="black")

#abline(v=tst.t[store.R0s1[,2]==max(store.R0s1[,2])])

points(tst.t[store.R0s1[,2]==max(store.R0s1[,2])],max(store.R0s1[,2]),pch=19,cex=1.2,col="grey")

\## For parameter r1 ##########################

\### 1. for low mortality

chs.k <- 30

tst.r <- seq(0,0.4,length=100)

store.R0s <- matrix(NA,length(tst.r),2)

for (jx in 1:length(tst.t)) {

store.R0s[jx,1] <- R0(Lt=5,k=aa$ktest[chs.k],d=0,t=aa$storePar[chs.k,1],v=1,r1=tst.r[jx],

r2=min(aa$storePar[chs.k,3],1-tst.r[jx]),r3=max(1-sum(aa$storePar[chs.k,3]+tst.r[jx]),0), mu1=0.01)

store.R0s[jx,2] <- R0hemi(Lt=5,k=aa$ktest[chs.k],d=0,t=aa$storePar1[chs.k,1],v=1,r1=tst.r[jx],r2=1-tst.r[jx],r3=0, mu1=0.01)

}

\### 2. for high mortality

store.R0s1 <- matrix(NA,length(tst.r),2)

for (jx in 1:length(tst.t)) {

store.R0s1[jx,1] <- R0(Lt=5,k=aa1$ktest[chs.k],d=0,t=aa1$storePar[chs.k,1],v=1,r1=tst.r[jx],

r2=min(aa1$storePar[chs.k,3],1-tst.r[jx]),r3=max(1-sum(aa1$storePar[chs.k,3]+tst.r[jx]),0), mu1=0.5)

store.R0s1[jx,2] <- R0hemi(Lt=5,k=aa1$ktest[chs.k],d=0,t=aa1$storePar1[chs.k,1],v=1,r1=tst.r[jx],r2=1-tst.r[jx],r3=0, mu1=0.5)

}

par(mfrow=c(1,1),bty="l")

plot(tst.r, store.R0s[,1], type="l", xlab=expression(r[1]), ylab=expression(R[0]), ylim=range(store.R0s,c(store.R0s1),na.rm=TRUE),lwd=1.5)

points(tst.r,store.R0s[,2], type="l", col="grey",lty=3,lwd=1.5)

#abline(v=tst.r[store.R0s[,1]==max(store.R0s[,1])])

points(tst.r[store.R0s[,1]==max(store.R0s[,1])],max(store.R0s[,1]),pch=19,cex=1.2,col="black")

#abline(v=tst.r[store.R0s[,2]==max(store.R0s[,2])])

points(tst.r[store.R0s[,2]==max(store.R0s[,2])],max(store.R0s[,2]),pch=19,cex=1.2,col="grey")

#plot(tst.r, store.R0s1[,1], type="l", xlab="t", ylab=expression(R[0]), ylim=range(store.R0s1,na.rm=TRUE),lwd=1.5)

points(tst.r,store.R0s1[,1], type="l", col="black",lty=1,lwd=1.5)

points(tst.r,store.R0s1[,2], type="l", col="grey",lty=3,lwd=1.5)

#abline(v=tst.r[store.R0s1[,1]==max(store.R0s1[,1])])

points(tst.r[store.R0s1[,1]==max(store.R0s1[,1])],max(store.R0s1[,1]),pch=19,cex=1.2,col="black")

#abline(v=tst.r[store.R0s1[,2]==max(store.R0s1[,2])])

points(tst.r[store.R0s1[,2]==max(store.R0s1[,2])],max(store.R0s1[,2]),pch=19,cex=1.2,col="grey")

\## For r2 ##########################

\### 1. for low mortality

chs.k <- 30

tst.r <- seq(0,0.9,length=100)

store.R0s <- matrix(NA,length(tst.r),2)

for (jx in 1:length(tst.t)) {

store.R0s[jx,1] <- R0(Lt=5,k=aa$ktest[chs.k],d=0,t=aa$storePar[chs.k,1],v=1,r2=tst.r[jx],

r1=min(aa$storePar[chs.k,2],1-tst.r[jx]),r3=max(1-sum(aa$storePar[chs.k,2]+tst.r[jx]),0), mu1=0.01)

store.R0s[jx,2] <- R0hemi(Lt=5,k=aa$ktest[chs.k],d=0,t=aa$storePar1[chs.k,1],v=1,r2=tst.r[jx],r1=1-tst.r[jx],r3=0, mu1=0.01)

}

\### 2. for high mortality

store.R0s1 <- matrix(NA,length(tst.r),2)

for (jx in 1:length(tst.t)) {

store.R0s1[jx,1] <- R0(Lt=5,k=aa1$ktest[chs.k],d=0,t=aa1$storePar[chs.k,1],v=1,r2=tst.r[jx],

r1=min(aa1$storePar[chs.k,2],1-tst.r[jx]),r3=max(1-sum(aa1$storePar[chs.k,2]+tst.r[jx]),0), mu1=0.5)

store.R0s1[jx,2] <- R0hemi(Lt=5,k=aa1$ktest[chs.k],d=0,t=aa1$storePar1[chs.k,1],v=1,r2=tst.r[jx],r1=1-tst.r[jx],r3=0, mu1=0.5)

}

par(mfrow=c(1,1),bty="l")

plot(tst.r, store.R0s[,1], type="l", xlab=expression(r[1]), ylab=expression(R[0]), ylim=range(store.R0s,c(store.R0s1),na.rm=TRUE),lwd=1.5)

points(tst.r,store.R0s[,2], type="l", col="grey",lty=3,lwd=1.5)

#abline(v=tst.r[store.R0s[,1]==max(store.R0s[,1])])

points(tst.r[store.R0s[,1]==max(store.R0s[,1])],max(store.R0s[,1]),pch=19,cex=1.2,col="black")

#abline(v=tst.r[store.R0s[,2]==max(store.R0s[,2])])

points(tst.r[store.R0s[,2]==max(store.R0s[,2])],max(store.R0s[,2]),pch=19,cex=1.2,col="grey")

#plot(tst.r, store.R0s1[,1], type="l", xlab="t", ylab=expression(R[0]), ylim=range(store.R0s1,na.rm=TRUE),lwd=1.5)

points(tst.r,store.R0s1[,1], type="l", col="black",lty=1,lwd=1.5)

points(tst.r,store.R0s1[,2], type="l", col="grey",lty=3,lwd=1.5)

#abline(v=tst.r[store.R0s1[,1]==max(store.R0s1[,1])])

points(tst.r[store.R0s1[,1]==max(store.R0s1[,1])],max(store.R0s1[,1]),pch=19,cex=1.2,col="black")

#abline(v=tst.r[store.R0s1[,2]==max(store.R0s1[,2])])

points(tst.r[store.R0s1[,2]==max(store.R0s1[,2])],max(store.R0s1[,2]),pch=19,cex=1.2,col="grey")

###############################################################################################################################################################################

\### Loop over d and mu1 to identify the switch point (Lt has not effect) ####

ntest <- 200

d.test <- seq(0,0.1,length=2)

mu1.test <- seq(0.01,0.5,length=100)

store.rc <- array(dim=c(length(d.test),length(mu1.test)))

store.gradient <- array(dim=c(length(d.test),length(mu1.test)))

par.homo <- array(dim=c(length(d.test),length(mu1.test),ntest,3))

par.hemi <- array(dim=c(length(d.test),length(mu1.test),ntest,3))

holo.hemi.distance <- array(dim=c(length(d.test),length(mu1.test)))

holo.hemi.params.dist.lowgrow <- holo.hemi.params.dist.highgrow <- holo.hemi.params.dist <- array(dim=c(length(d.test),length(mu1.test),3))

switch.index <- array(dim=c(length(d.test),length(mu1.test)))

#for (j in 1:length(d.test)) {

j<-1

for (k in 1:length(mu1.test)) {

tpp <- findKswitch(d=d.test[j],mu1=mu1.test[k],R=1,Lt=5, ntest=ntest,nest=40)

store.rc[j,k] <- tpp$switch.point

store.gradient[j,k] <- tpp$kgradient

par.homo[j,k,,] <- tpp$storePar

par.hemi[j,k,,] <- tpp$storePar1

switch.index[j,k] <- tpp$switch.index

before.switch <- max(switch.index[j,k]-1,1) #switch.index[j,k]#

if (length(before.switch)>0) {

holo.hemi.distance[j,k] <- sum((par.homo[j,k,before.switch,c(1,2,3)]-par.hemi[j,k,before.switch,c(1,2,3)])^2)

holo.hemi.params.dist[j,k,] <- par.hemi[j,k,before.switch,c(1,2,3)]-par.homo[j,k,before.switch,c(1,2,3)]

holo.hemi.params.dist.lowgrow[j,k,] <- par.hemi[j,k,1,c(1,2,3)]-par.homo[j,k,1,c(1,2,3)]

holo.hemi.params.dist.highgrow[j,k,] <- par.hemi[j,k,ntest,c(1,2,3)]-par.homo[j,k,ntest,c(1,2,3)]

}

}#}

cols <- colorRampPalette(c("blue", "red"))(ntest)

\## just look at death for d=0 (d=1 is consistent pattern)

par(mfrow=c(1,1), bty="l")

plot(mu1.test, store.rc[1,], type="l",

xlab=expression("Mortality during growth phase, "*mu[G]),

ylab=expression("Threshold "*italic(k)*" for holometab."),lwd=1.3)

\## plot with full range

par(mfrow=c(1,1), bty="l")

plot(mu1.test, store.rc[1,], type="l",

xlab=expression("Mortality during growth phase, "*mu[G]),

ylab=expression("Threshold "*italic(k)*" for holometab."),lwd=1.3, ylim=c(0,1))

\## just look at death for d=0 - at a particular k (not the switch one)

par(mfrow=c(3,4), bty="l", mar=c(5,4,1,1))

for (kchs in c(1,10,20)) {

plot(mu1.test, par.homo[1,,kchs,1], type="l",

xlab=expression("Mortality during growth phase, "*expression(mu[G])),

ylab=expression(italic(t)),lwd=1.3)

points(mu1.test, par.hemi[1,,kchs,1], type="l",lty=3,col="grey")

plot(mu1.test, par.homo[1,,kchs,2], type="l",ylim=c(0,1),

xlab=expression("Mortality during growth phase, "*expression(mu[G])),

ylab=expression(italic(r[1])),lwd=1.3)

points(mu1.test, par.hemi[1,,kchs,2], type="l",lty=3,col="grey")

plot(mu1.test, par.homo[1,,kchs,3], type="l",ylim=c(0,1),

xlab=expression("Mortality during growth phase, "*expression(mu[G])),

ylab=expression(italic(r[2])),lwd=1.3)

points(mu1.test, par.hemi[1,,kchs,3], type="l",lty=3,col="grey")

plot(mu1.test, 1-par.homo[1,,kchs,2]-par.homo[1,,kchs,3], type="l",ylim=c(0,1),

xlab=expression("Mortality during growth phase, "*expression(mu[G])),

ylab=expression(italic(r[3])),lwd=1.3)

}
